# Combinatorial Polyacrylamide Hydrogels for Preventing Biofouling on Implantable Biosensors

**DOI:** 10.1101/2020.05.25.115675

**Authors:** Doreen Chan, Jun-Chau Chien, Eneko Axpe, Louis Blankemeier, Samuel W. Baker, Sarath Swaminathan, Victoria A. Piunova, Dmitry Yu. Zubarev, Caitlin L. Maikawa, Abigail K. Grosskopf, Joseph L. Mann, H. Tom Soh, Eric A. Appel

**Affiliations:** Department of Chemistry, Stanford University, Stanford, CA 94305, USA; Department of Materials Science & Engineering, Stanford University, Stanford, CA 94305, USA; Department of Electrical Engineering, Stanford University, Stanford, CA 94305, USA; Department of Comparative Medicine, Stanford University, Stanford, CA 94305, USA; IBM Almaden Research Center San Jose, CA 95120, USA; Department of Bioengineering, Stanford University, Stanford, CA 94305, USA; Department of Radiology, Stanford University, Stanford, CA 94305, USA; Chan Zuckerberg Biohub, San Francisco, CA 94158, USA; ChEM-H Institute, Stanford University, Stanford, CA 94305, USA; Department of Pediatrics (Endocrinology), Stanford University, Stanford, CA 94305, USA

**Keywords:** anti-fouling, polymers, biosensors, devices, hydrogels

## Abstract

Biofouling on the surface of implanted medical devices severely hinders device functionality and drastically shortens device lifetime. Poly(ethylene glycol) and zwitterionic polymers are currently considered “gold standard” device coatings to reduce biofouling. To discover novel anti-biofouling materials, we created a combinatorial library of polyacrylamide-based copolymer hydrogels and screened their ability to prevent fouling from serum and platelet-rich plasma in a high-throughput parallel assay. We found certain non-intuitive copolymer compositions exhibit superior antibiofouling properties over current gold standard materials, and employed machine learning to identify key molecular features underpinning their performance. For validation, we coated the surfaces of electrochemical biosensors with our hydrogels and evaluated their anti-biofouling performance *in vitro* and *in vivo* in rodent models. Our copolymer hydrogels preserved device function and enabled continuous measurements of a small-molecule drug *in vivo* better than gold standard coatings. The novel methodology we describe enables the discovery of anti-biofouling materials that can extend the lifetime of real-time *in vivo* sensing devices.

## INTRODUCTION

Medical devices improve the quality of life and extend the lifespan of millions of people worldwide^1^. But after implantation in the body of a patient, biofouling—the non-specific adsorption of biomolecules—often occludes these devices and contributes to a reduction in their performance and lifetime. For devices that contact blood, the adhesion and aggregation of platelets from flowing blood significantly shorten functional device lifetimes^2,3^. Ventricular devices, intravenous cannulas, and stents are all at risk of causing thrombosis, which can increase risk of embolism and stroke^4^. Proteins can adhere to indwelling central venous catheters within 24 hours, and the resulting thrombosis can lead to pain and infection^5,6^. Once these devices foul, high-risk and costly invasive surgeries are often required to replace them, greatly increasing patient and hospital burden.

To address these challenges, devices are often coated with a soft material to mitigate biofouling. “Active” coatings such as drug-eluting^7^ or lubricant-infused polymer materials^8^ can be used to reduce adhesion and activation of platelets. While these approaches show promise in some applications, they offer only short-term benefits on account of the eventual depletion of the active molecules from the coating material^9–11^, leading to major bleeding in patients near the end of the functional device lifetime^12^. In contrast, “passive” coating strategies that modify the chemistry at the interface of the devices and bodily fluids are scalable, tunable, and inexpensive. The “gold standard” materials used for passive anti-biofouling coatings are poly(ethylene glycol) (PEG)^13^ and its derivatives^14–16^, which form a tight hydration layer through hydrogen bonding with water that is hypothesized to contribute to anti-biofouling properties^17^. Nonetheless, PEG has several critical drawbacks, including degradation through hydrolysis and auto-oxidation leading to the production of reactive oxygen species^18–21^ and reduced anti-fouling performance over time^22^. In recent years, many alternatives to PEG have been developed^23^, including zwitterionic polymers^24,25^, fluoro- and acrylate-based polymers^26–28^, and polyglycidols^29^. Poly-zwitterionic materials in particular have shown promise as exceptional anti-biofouling coatings, though these materials may still face reduced stability in long-term applications due to the presence of hydrolysis-prone ester bonds^30–32^. There is thus a critical need to develop materials that achieve superior stability and anti-biofouling performance relative to existing materials. Novel approaches that make use of high-performance assays to rigorously test the function of materials are needed to advance the discovery of such next-generation anti-biofouling materials, enabling the development of coatings that improve the performance of medical devices.

In this study, we used a high-throughput screening approach to evaluate biofouling of a library of 172 combinatorial copolymer hydrogel materials assembled from 11 distinct acrylamide-based monomers (**Fig. 1**). Our results demonstrate that a subset of these materials exhibit remarkable anti-biofouling function compared to existing gold standard coatings. These polyacrylamide hydrogels exhibit tunable mechanical properties, and the simplicity of their synthesis makes them highly adaptable for biosensor coatings. Although combinatorial screening approaches have been employed previously for materials discovery efforts^33–35^ aimed at mitigating marine or bacterial fouling^36–39^, the foreign body response^35,40^, or cell adhesion^41,42^, to the best of our knowledge, this is the first work to screen for the prevention of platelet adhesion, which is a critical initial process for biofouling in blood. We have rigorously assessed the anti-fouling behavior of our library of copolymer hydrogels using a high concentration of serum proteins and platelet-rich plasma over prolonged timeframes. Moreover, we have used machine learning techniques to elucidate the molecular features of these combinatorial copolymer hydrogels that give rise to their observed biofouling performance. We show that our top-performing copolymer hydrogel is superior to current gold standard materials in terms of enhancing the performance and lifetime of electrochemical biosensors *in vitro* and *in vivo* in multiple rodent models.

**Figure 1.**
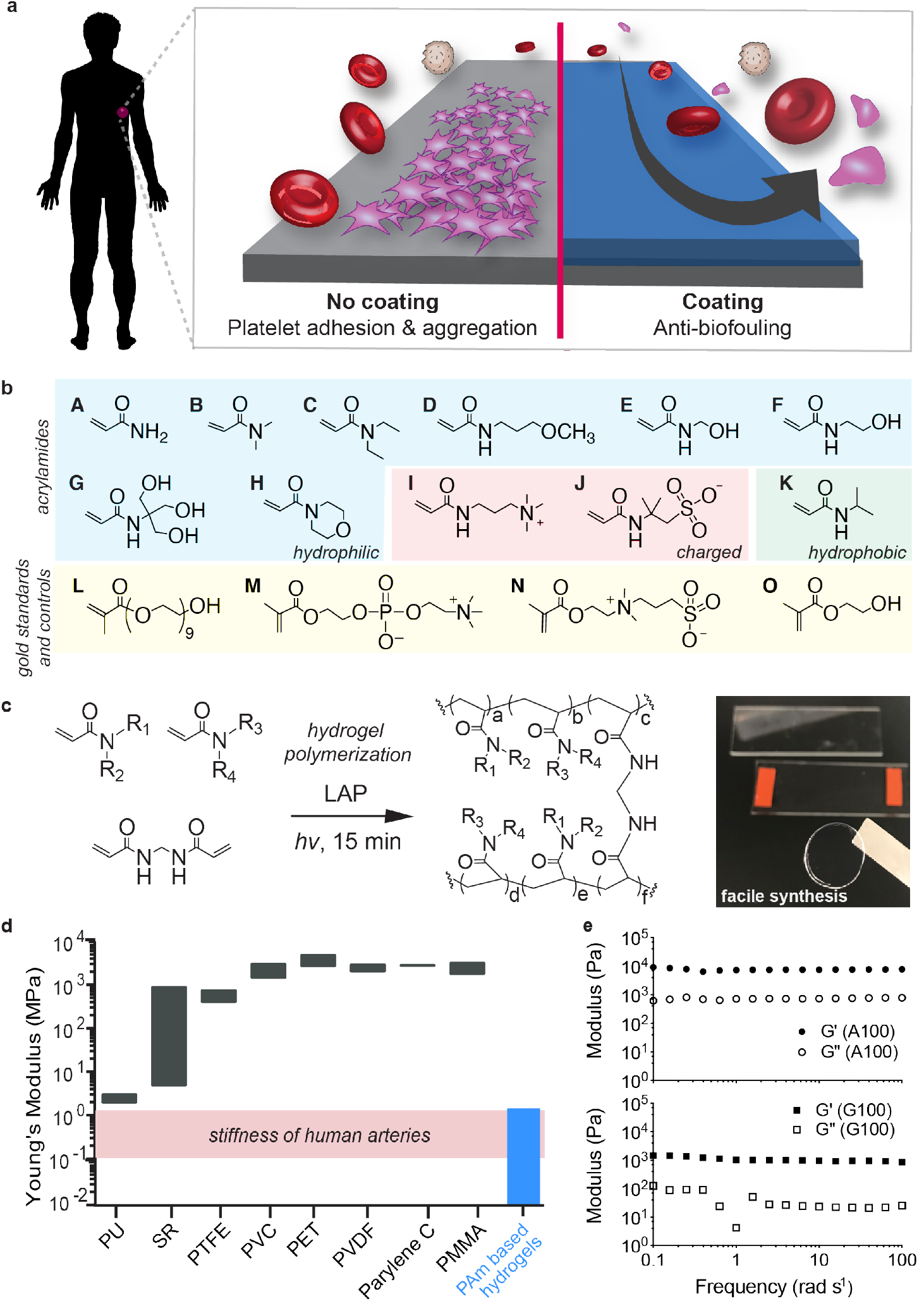
Polyacrylamide hydrogels as anti-biofouling device coatings. **a.** Coating substrates with anti-biofouling hydrogels can mitigate platelet adhesion and aggregation. **b.** Monomers employed for combinatorial hydrogel synthesis: acrylamide (A), dimethylacrylamide (B), diethylacrylamide (C), (3-methoxypropyl)acrylamide (D), hydroxymethylacrylamide (E), hydroxyethylacrylamide (F), [tris(hydroxymethyl)methyl]acrylamide (G), acryloylmorpholine (H), (acrylamidopropyl)trimethyl-ammonium (I), 2-acrylamido-2-methyl-propane sulfonic acid (J), N-isopropylacrylamide (K), poly(ethylene glycol) methacrylate (L), 2-methacryloyloxyethyl phosphorylcholine (M), [2- (methacryloyloxy)ethyl]dimethyl(3-sulfopropyl)ammonium (N), 2-hydroxyethyl methacrylate (HEMA; O). **c.** Photopolymerization (λ = 350 nm) of polyacrylamide hydrogels using lithium phenyl-2,4,6-trimethylbenzoylphosphinate (LAP) as a radical photoinitiator. **d.** Range of Young’s moduli for commercial polymers: polyurethane (PU), silicone rubbers (SR), polytetrafluoroethylene (PTFE), polyvinyl chloride (PVC), polyethylene terephthalate (PET), parylene C, and poly(methyl methacrylate) (PMMA). In this study, polyacrylamide copolymer hydrogels were fabricated with elastic moduli similar to human arteries. **e.** Oscillatory shear rheology of hydrogels formed with low (A100; top) and high (G100; bottom) molecular weight monomers indicates consistency in shear moduli amongst the hydrogels.

### Design of a Combinatorial Library of Polyacrylamide Hydrogels for Anti-biofouling

Here we first sought to investigate the ability of a large library of chemically distinct polyacrylamide copolymer hydrogels to mitigate blood protein and platelet adhesion (**Fig. 1a**). Several acrylamide-derived monomers have been widely utilized in biological settings (*e.g*., SDS-PAGE gels) and in biomedical applications (*e.g*., contact lenses, fillers, and drug delivery materials) and have previously demonstrated potential for use as inert materials in the human body^43,44^. To maximize the scope of our evaluation of polyacrylamide materials, we selected 11 commercially-available acrylamide-derived monomers and fabricated a library of 172 polyacrylamide copolymer hydrogels comprising unique binary combinatorial mixtures (100:0, 75:25, 50:50, 25:75) of these monomers formulated at 20 wt% monomer (**Fig. 1b, 1c; Supplementary Tables 1 and 2**). To the best of our knowledge, this library represents the largest combinatorial library of polyacrylamide-based hydrogel formulations evaluated to date.

Each of the hydrogels were prepared by photopolymerization of prepolymer solutions using lithium phenyl-2,4,6-trimethylbenzoylphosphinate (LAP) as a radical photoinitiator and a simple LED (λ = 350 nm) light source. During synthesis, four copolymer formulations turned opaque on account of insolubility of the polymer chains in the aqueous media and were thus omitted from further evaluation. We synthesized our hydrogels at a monomer content of 20 wt% to generate a library of materials with stiffness values mimicking those of human vein or artery tissues (**Fig. 1d**)^45,46^. As all monomers evaluated were acrylamide-derived, the reactivity ratios for copolymerization of each pair of monomers (r1r2 ≈ 1) were such that each copolymer hydrogel was expected to comprise a statistical incorporation of each monomer moiety throughout the hydrogel network^47^. We further sought to ensure that subsequent assays would strictly screen for the impact of polymer chemistry on anti-biofouling performance, rather than mechanics. We performed oscillatory shear rheology on the homopolymer hydrogels comprising the two monomers with the most distinct molecular weights: acrylamide (A; M_w_ = 71.08 g/mol) and [tris(hydroxymethyl)methyl]-acrylamide (G; M_w_ = 175.18 g/mol). We selected these two formulations for more in-depth mechanical characterization in order to validate that the various members of our hydrogel library exhibited similar mechanical properties (**Fig. 1e**). These studies indicated that even these highly distinct hydrogels exhibited similar shear storage and loss modulus values, and elastic modulus values similar to human veins and arteries.

### Protein Adsorption and Platelet-Resistant Hydrogels

To rapidly evaluate the extent of biofouling on these hydrogels in a parallel fashion, we developed a high-throughput assay to screen platelet adhesion following incubation in protein serum. While the exact mechanism of biofouling has yet to be precisely elucidated, it is widely hypothesized that such fouling is initiated with the non-specific adsorption of proteins from the biological milieu^48^. This undesirable binding of proteins to the surface of a biomaterial or device leads to the onset of platelet adhesion and aggregation that eventually causes thrombosis^48–51^. While the majority of anti-biofouling assays appearing in the literature have focused on evaluating adsorption of model proteins at low concentrations (*e.g*., 1 mg/mL bovine serum albumin^39^ or 1 mg/mL fibrinogen^52^) onto materials of interest, these studies do not provide a realistic representation of the complex milieu of biomolecules in blood^53^. Other studies have utilized more relevant mixtures of biological proteins (*e.g*., fibrinogen, bovine serum albumin, and lysozyme), or diluted or undiluted serum or plasma to evaluate biofouling of new materials, but only over short exposure timeframes ranging from 10–25 minutes^54,24^. The ability to repel whole blood has also been studied; however, in these studies blood was applied to the biomaterial of interest for only a matter of seconds^55^. Consequently, previous studies have not provided a sufficiently realistic representation of the complex engineering challenge imposed on anti-biofouling coatings in the body on account of low protein concentrations, insufficiently complex biological milieu, and short assay times^53^.

To identify anti-biofouling materials from our copolymer hydrogel library with translational potential, we therefore sought to subject these materials to severe fouling conditions for prolonged periods of time that would nevertheless provide a simple and high-throughput readout of fouling activity. We chose to use platelet counting to enable a straightforward and realistic metric for anti-biofouling performance of our copolymer hydrogels, as platelet adhesion causes occlusion and leads to thrombosis^7^. In these assays, each hydrogel was first incubated in 50% fetal bovine serum (FBS) for 24 hr at 37 °C to introduce non-specific protein adsorption (**Fig. 2a**) prior to incubation in platelet-rich plasma (PRP) for 1 hr at 37°C to introduce platelets that could be counted in an automated fashion. These time-points were chosen to allow for discrimination between the top formulations while minimizing deviation as greater platelet counts introduced higher variation (**Fig. 2b**). Although we could not discern clear trends in the behaviors of the polyacrylamide copolymers based on their composition, hydrophobic monomers such as N-isopropylacrylamide (K100) or diethylacrylamide (C100) yielded high mean platelet counts (28,600 ± 11,000 platelets/cm^2^ and 9,870 ± 13 platelets/cm^2^, respectively). In contrast, a 50:50 copolymer of hydroxyethylacrylamide (F) and diethylacrylamide (C), denoted F50-C50, yielded the lowest platelet counts (82.0 ± 15 platelets/cm^2^) of all of the materials tested, significantly outperforming PEG (L; 4,810 ± 520 platelets/cm^2^; p < 0.0001), 2-methacryloyloxyethyl phosphorylcholine (M; 1,330 ±210 platelets/cm^2;^ p = 0.0001), poly(2-hydroxyethyl methacrylate) (PHEMA; O; 9,560 ± 650 platelets/cm^2^; p < 0.0001), and [(methacryloyloxy)ethyl]-dimethyl(3-sulfopropyl)ammonium zwitterion (N; 15,900 ± 4,200 platelets/cm^2^; p = 0.0007) hydrogel formulations (mean ± standard error, unpaired *t* test) (**Fig. 2c, Supplementary Fig. 1**). This approach to high-throughput screening of fouling with blood proteins and platelets therefore enabled identification of polyacrylamide-based copolymer hydrogel formulations with superior anti-biofouling properties in comparison to gold standard polymers. While protein adsorption has previously been investigated as a metric of anti-biofouling performance, we did not see trends correlating IgG and fibrinogen adsorption with platelet adhesion (**Supplementary Fig. 2**). This observation may indicate that other proteins and/or specific conformations of adsorbed proteins (rather than quantity alone) may play key roles in platelet adhesion. As protein layers evolve unpredictably over time, protein adsorption may offer limited utility as a metric correlating to platelet adhesion^56^.

**Figure 2.**
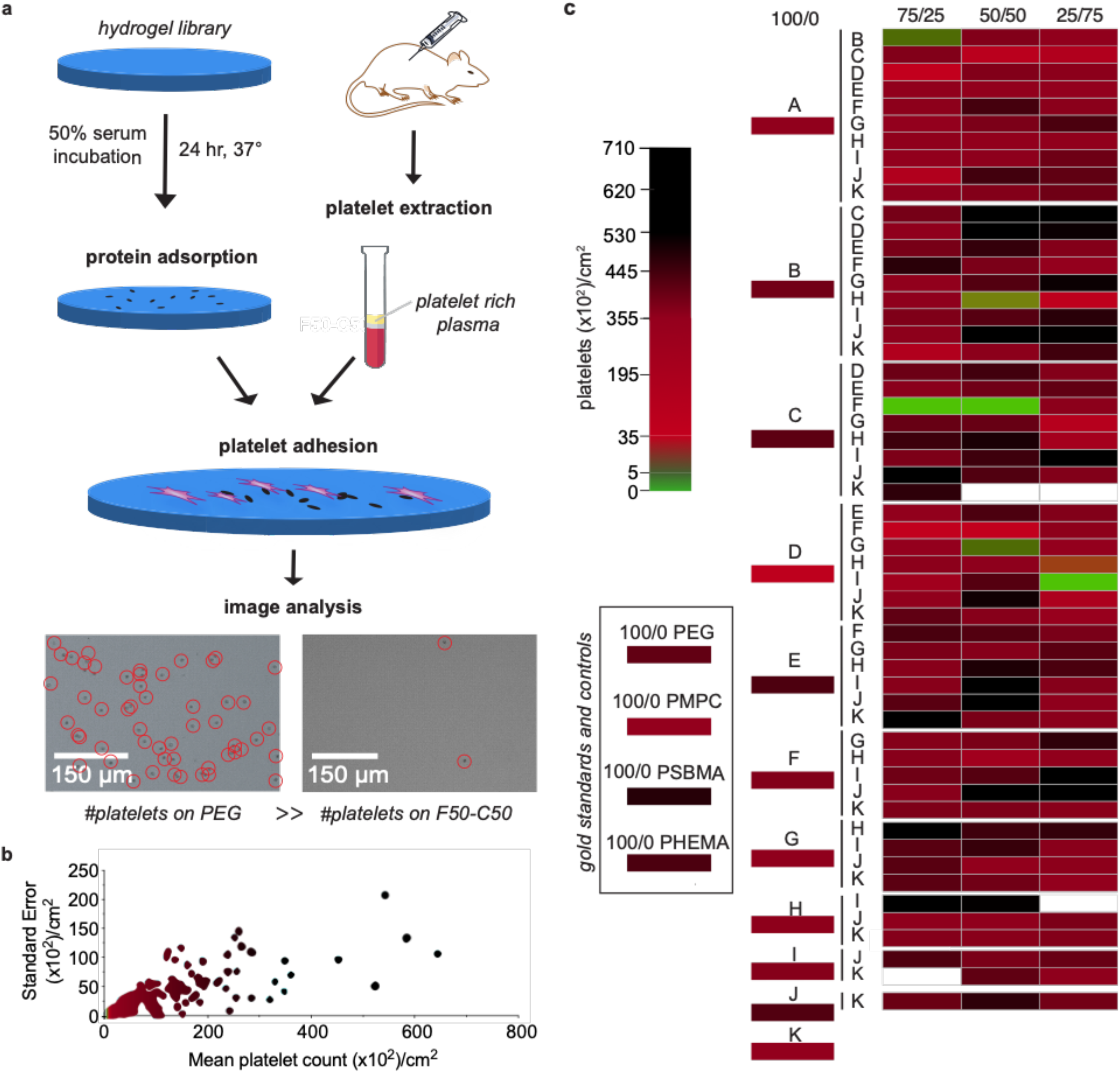
Identification of anti-biofouling acrylamide-based copolymer combinations. **a.** Hydrogel samples were first incubated in 50% fetal bovine serum for 24 hr at 37 °C to ensure extensive protein adsorption to the samples. Platelet rich plasma (PRP) obtained from centrifugation of rat blood was then incubated on the surface of the hydrogels for 1 hr at 37 °C. Several polyacrylamide copolymer hydrogels presented significantly less platelet adhesion than PEG, PHEMA, and polyzwitterionic hydrogel controls. **b.** Standard error versus mean of the platelet count distribution obtained for each copolymer hydrogel. **c.** Heat map of median platelet adhesion counts on copolymer hydrogel samples, ordered by the weight ratio of each monomer in the hydrogel formulation (100/0, 25/75, 50/50, 75/25). Colors were assigned to the medians obtained in n ≥ 3 tests.

We then performed scanning electron microscopy (SEM) imaging of treated hydrogel samples to observe platelet morphology as an indicator of platelet activation. On the PEG hydrogel sample, aggregates of platelets were observed throughout the surface of the hydrogel, although their morphology was generally round, indicating that the platelets were not activated^57^. Similarly, the PHEMA hydrogel exhibited aggregates of round platelets present across its surface^56^. In contrast, polyzwitterionic materials contained fewer platelet aggregates, but the morphology of these platelets was indicative of activation—we observed that these platelets had morphed from spheroid form to exhibit bulbous protrusions on their surface. On the F50-C50 polyacrylamide hydrogels, we observed negligible platelet aggregation across the surface and little to no platelet activation (**Supplementary Fig. 3**).

### Identifying Key Features of Monomers for Anti-Biofouling Properties

To aid in our understanding of the molecular features giving rise to the observed anti-biofouling properties of all of the materials we tested, we analyzed the physicochemical properties of each of the monomers and resulting hydrogels and the contribution of these features to fouling tendencies^58^. The use of platelet counts allowed for inputs that can link polymer behaviors to antibiofouling performance, in contrast to protein adsorption assays that may not yield predictive models^41^. The long-standing hypothesis regarding the anti-fouling mechanism of the gold standard polymers is the presence of a tight hydration layer between the material surface and water that forms a physical and energetic barrier to protein adsorption, determined often by the physicochemical properties of the materials and the surface packing^59^. These properties include surface wettability and hydrophilicity, electrical neutrality, H-bonding, and prevalence of highly hydrated chemical groups. It has also been suggested that surface packing has a role in non-fouling properties, strongly correlated with the ability to form a hydration layer near the surface. Molecular simulations have also suggested that anti-biofouling properties may arise from steric repulsion of proteins and polymer surfaces - that the density of grafted polymer chains simply causes resistance to protein adhesion and obstructs their adsorption to the surface^17,60–62^. Numerous other computational and molecular dynamics studies have shown that entropic contributions play a role in preventing biofouling^63,64^, wherein reduction in ligand mobility would lead to an entropic penalty against protein adsorption^65^. Penna *et al*. have identified the importance of ligand mobility and interfacial dynamics, highlighting a balance of hydration properties and a dynamic ligand layer that greatly affects entropic barriers associated with antifouling efficacy^66^.

Curiously, the compositions of the top-performing hydrogel formulations that yielded the lowest degree of platelet adhesion did not exhibit any intuitive trends, and so we utilized machinelearning analysis to identify mechanistic factors underpinning anti-biofouling activity. We first classified the copolymer hydrogels as “low platelet count” or “high platelet count” with a random forest model, which offers robustness to overfitting and the capability to evaluate relevance of data features via mean decrease of accuracy (MDA) (see Methods for technical details). We initially generated a simple feature set (Feature Set A) reflective of the polymer formulation, whereby each polymer was represented using one-hot encoding for the alphabet of monomers weighted by the molar ratios of the monomers in the respective copolymer hydrogels. Feature relevance analysis in combination with class-enrichment analysis (**Fig. 3a, Supplementary Fig. 4a**) pointed to dimethylacrylamide (B), diethylacrylamide (C), and (acrylamidopropyl)trimethylammonium (I) being associated with heavy biofouling. In contrast, acrylamide (A), (3-methoxypropyl)acrylamide (D), and acryloylmorpholine (H) were prominent monomers in those hydrogel formulations exhibiting excellent anti-biofouling performance. As none of these monomers were in the top anti-biofouling hydrogel formulations, we determined that combinatorial formulations, rather than homopolymer formulations, are crucial in development of robust anti-biofouling coatings.

**Figure 3.**
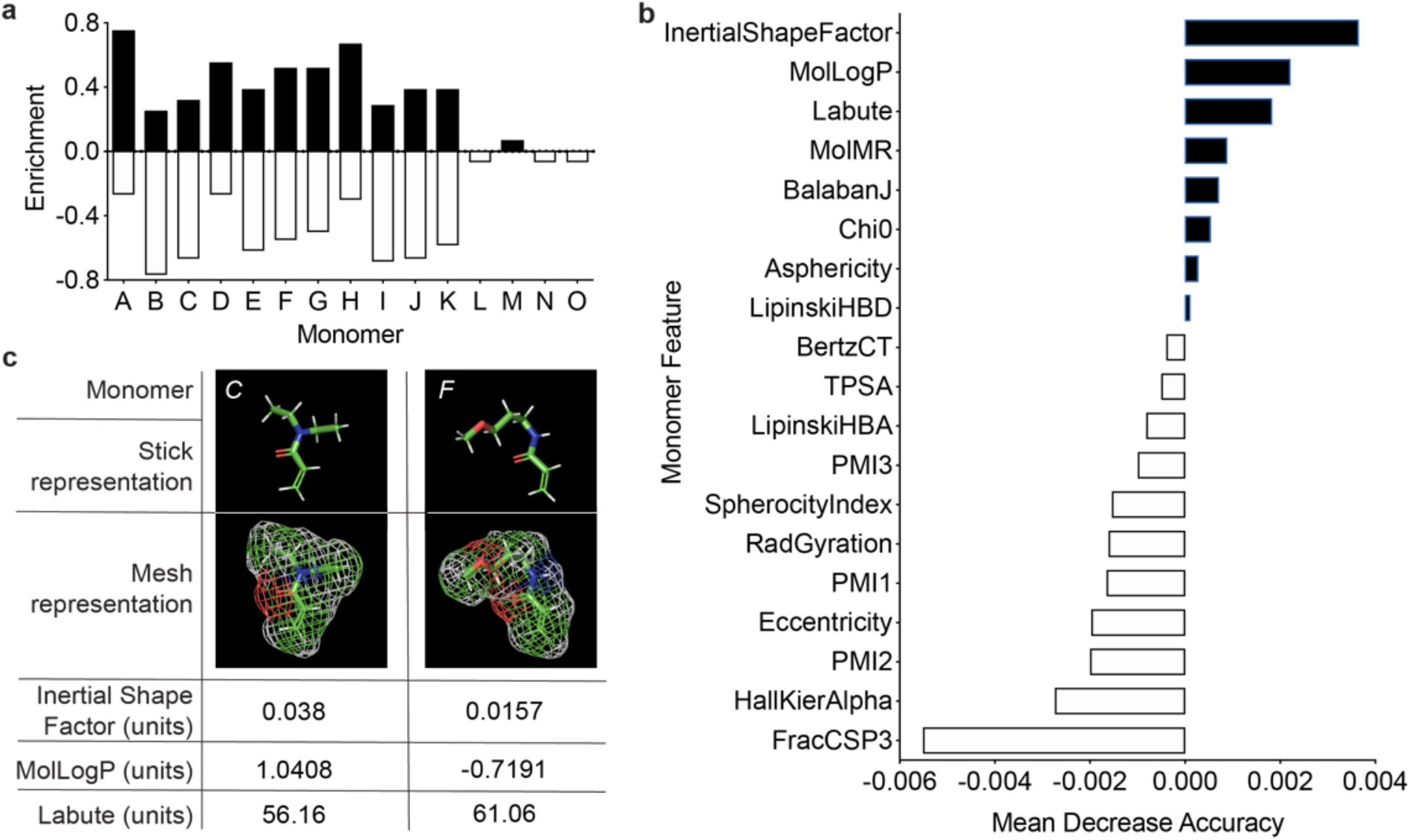
Feature relevance evaluated for monomers using trained random forest models. **a.** Monomer-specific class enrichment based on raw platelet adhesion data, indicating contribution of individual monomers to anti-biofouling performance. This does not account for the inherent features of the monomers, but simply quantifies the contribution of each specific monomer. **b.** Mean decrease of accuracy (MDA) of descriptor values evaluated with Feature Set B, summing the importance of the features across all monomers. Positive values (black) indicate that the feature is relevant to model performance, while negative (white) values do not inform the model. **c.** Descriptor values with the highest overall MDA for monomers in the copolymer hydrogel with the best anti-biofouling performance (F50-C50). Descriptors are obtained by transforming complex chemical information about the monomer into a number suitable for analysis.

This consideration motivated two approaches of weighing both the molecular features of each specific monomer and the ratio of each monomer present in a specific formulation. In this feature set, we increased resolution of the molecular features by generating vectors of physicochemical descriptors for the monomers, “inserting” these vectors into one-hot monomer encoding, and weighting by the molar ratios of the monomers in the corresponding copolymer hydrogels (Feature Set B). The f1 score (the overall measure of a model’s accuracy) indicated that these feature sets were suitable in data analysis (**Supplementary Table 3**). Descriptors, in general, are numerical values that capture various characteristics of the molecules, both simple and complex. For example, a feature can be the number of proton donors, or the complexity measure of the molecular graph. The set of descriptors computed for each monomer included 19 values split between physicochemical descriptors (LabuteASA, TPSA, MolLogP, MolMR), molecular graph descriptors (BalabanJ, BertzCT, Chi0, HallKierAlpha), hydrogen-bonding descriptors (LipinskiHBA, LipinskiHBD), and 3D shape descriptors (FracCSP3, InertialShapeFactor, PMI1, PMI2, PMI3, RadGyration, SpherocityIndex, Asphericity, Eccentricity), all as implemented in the *rdkit* library^67^ (**Supplementary Table 4**).

Inspection of the feature analysis for the monomers weighted by their molar ratios helped to pinpoint relevant characteristics of the individual monomers. These features could then be expressed as descriptors (**Fig. 4b, Supplementary Table 4, Supplementary Fig. 5**). The top three features are from the physicochemical descriptor (MolLogP and LabuteASA) and 3D shape family (InertialShapeFactor). Overall, the most relevant features were found to be connectivity and shape descriptors: (i) InertialShapeFactor describes the mass distribution in the molecule (by three principal moments of inertia), (ii) MolLogP is an estimate of the octanol/water partition coefficient and captures the hydrophilicity/hydrophobicity of the molecule (an energetic characteristic), and (iii) LabuteASA describes the approximate surface area of the molecule. Surface area of the molecules plays a role in anti-biofouling performance because all monomers have a surface area that is accessible to other molecules (*e.g*. the solvent), and higher values of LabuteASA indicate that more surface is accessible (**Fig. 3c**).

**Figure 4.**
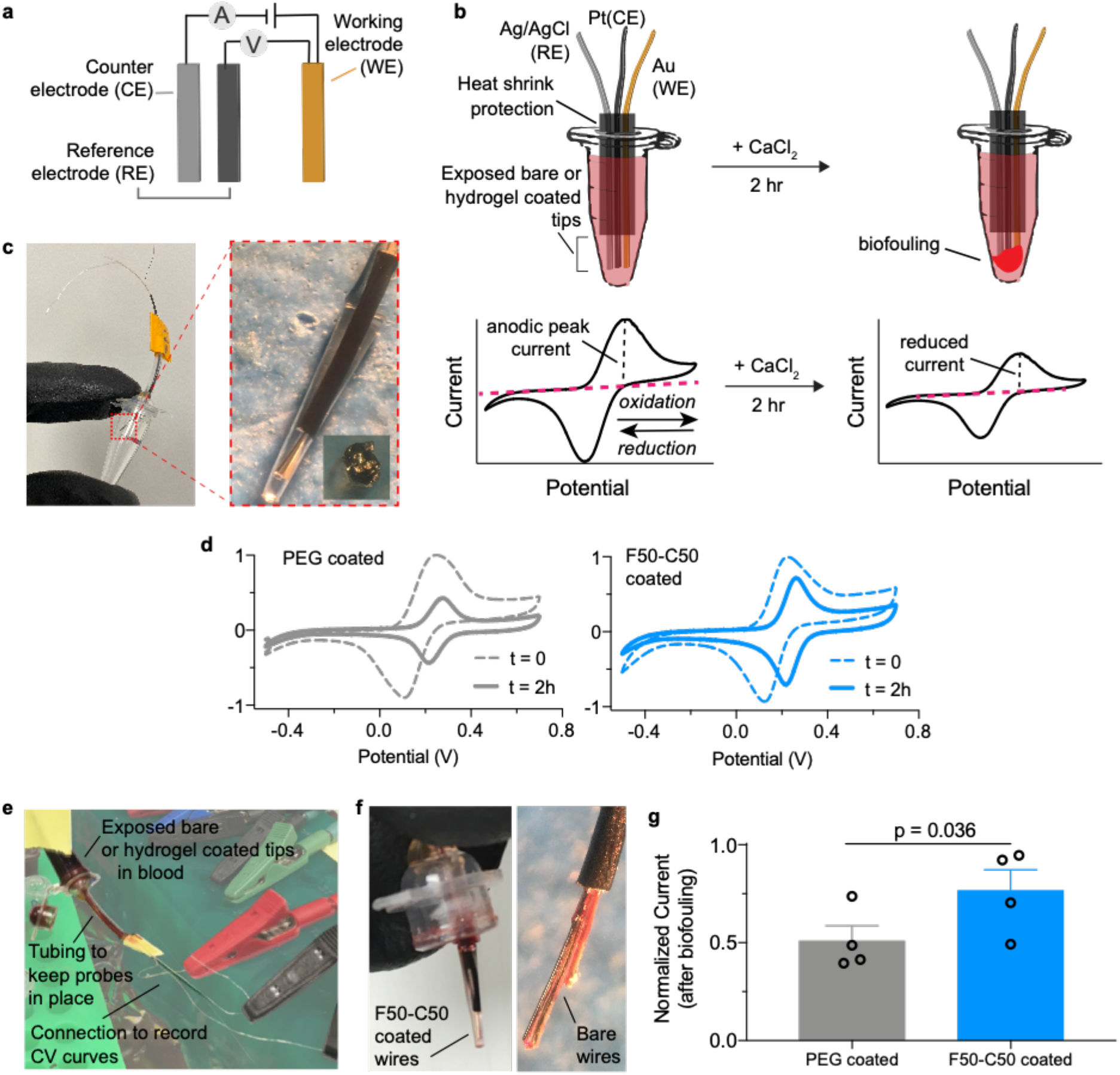
Hydrogel protection of an electrochemical device extends lifetime and signal quality over bare and PEG-coated devices. **a.** Affinity-based electrochemical sensors can be used to detect oxidation of ferrocyanide species (FeCN_6_)^4-^. **b.***In vitro* assay to observe blood fouling on the device surface. Probes are inserted through the Eppendorf tube cap to prevent contact with side walls. Biofouling on the device reduces cyclic voltammetry (CV) oxidation and reduction peaks. **c.** Probes were coated with a visible layer of hydrogel to mitigate biofouling. **d.** CV curves were obtained by cycling electrodes between oxidation and reduction potentials of ferrocyanide. Representative graphs are shown before and after CaCl_2_ addition. **e.***In vitro* assay setup. Tubes were inverted to ensure probes are completely immersed in blood. **f.** Devices show visible signs of fouling after 2 hr incubation. **g.** Normalized signal intensity, defined by anodic peak current from CV, of devices after biofouling. Data depict mean ± standard error, n =4. Significance was determined through a paired t-test.

In a second approach, we produced a simplified version of the descriptor-based feature set by summing up descriptor vectors of the monomers weighted by their molar ratios in corresponding copolymer hydrogels (Feature Set C), and our results indicated the importance of the 3D shape descriptor features (**Supplementary Fig. 4b**). Expanded analysis of the heterogeneity of the platelet count distribution also indicated the importance of a steric mode in the observed fouling (**Supplementary Fig. 6**). Overall, it appears that the relevant molecular features predominantly characterize properties associated with the steric aspect of the surface formation and the geometry of packing of the monomers.

### Mechanical and Physical Properties of the Top Performing Hydrogel

The mismatch in mechanical properties between many implant materials and native tissues can lead to adverse events such as immune rejection, fibrotic responses, trauma, and thrombosis^68^. Thus, it is crucial to match the mechanical properties of the biomaterial with surrounding tissue^69–71^. Hydrogels in particular are widely tunable, and have mechanical similarity to biological tissue – in particular the human artery and vein – making them useful for implanted biosensors^72^ (**Fig. 1d, 1e** and **Supplementary Fig. 7**). While other coating methods may incorporate polymers via either grafting-to or grafting-from approaches, these coatings are often only nanometers thick and have poor mechanical integrity, making them ill-suited to serve as a soft interface between implanted devices and the body.

We first assessed the mechanical properties of our top-performing hydrogel formulation and control hydrogels comprising PEG, poly(2-methacryloyloxyethyl phosphorylcholine) ( PMPC), poly(2-methacryloyloxyethyl phosphorylcholine) (PSBMA), and poly(2-hydroxyethyl methacrylate) (PHEMA) using rheology (**Supplementary Fig. 7, 8**). The shear rheological properties were comparable among each material. Moreover, we used fluorescence recovery after photobleaching (FRAP) spectroscopy to evaluate the mesh size of each of these hydrogel materials, which were found to be similar to each other and well below the size of albumin^73^ (**Supplementary Fig. 7f**). We also used interferometry-based nanoindentation, which enables examination of the mechanics of these materials under wet conditions (**Supplementary Fig. 8a, 8c**), as well as tensile testing (**Supplementary Fig. 8b, 8c**) to characterize the elastic modulus of our top-performing hydrogel formulation. While soft materials commonly used in the medical industry exhibit Young’s moduli values several orders of magnitude greater than that of human arteries, our F50-C50 hydrogel coating exhibits mechanical properties that are similar to human arteries^74^, reducing the risk of chronic inflammation and potential failure of the implant^75^. Further, polyacrylamide hydrogels offer improved stability under a variety of expedited degradation conditions relative to the control materials we evaluated (**Supplementary Fig. 9**).

### Hydrogel Coating of Electrochemical Biosensors Extends Sensor Lifetime

In order to assess the usefulness of our new copolymer hydrogel material, we tested its antibiofouling performance in protecting electrochemical biosensors in blood. Such biosensors generally offer a promising strategy for continuously detecting analytes *in vivo*, but their lifetimes are limited by biofouling^59,76^. The degree of biofouling can be quantified using cyclic voltammetry (CV) to measure the anodic peak current. By applying a range of voltages through an electrode, electron transfer and current intensity may be monitored through the anodic peak. Changes in this peak can be used to assess interaction between the electrode surface and (FeCN_6_)^4-^ ions in solution and the lifetime and quality of the signal. A larger reduction in signal indicates barriers to the electron transfer process and blocking of ions from diffusing toward the electrode surface due to biofouling^77^.

We prepared devices comprising a gold (Au) working electrode (WE), a silver/silver chloride (Ag/AgCl) reference electrode (RE), and a platinum (Pt) counter electrode (CE) (**Fig. 4a, 4b**) with PEG and F50-C50 hydrogel coatings to assess the ability of the hydrogel coating to extend device lifetime. We first performed CV measurements in Na_2_EDTA-treated human whole blood spiked with 15 mM of ferrocyanide ions, (FeCN_6_)^4-^, to establish a baseline (**Fig. 4c**). Then, devices were incubated in ferrocyanide-free human whole blood mixed with 50 mM CaCl_2_ to initiate expedited blood coagulation^78,79^ and to activate blood platelets (**Supplementary Fig. 10**). After 2 hr, probes were placed in the same ferrocyanide-spiked blood sample to record CV measurements (**Fig. 4d-f, Supplementary Fig. 11**). As the addition of CaCl_2_ rapidly expedites blood clotting, 2 hr was ample time to observe significant differences in probe performance without full clotting of the blood medium^78^. While anodic peak current was measured, the slight appearance of other peaks was observed with all probes, indicating a change in diffusion rate. The shift in potential often occurs due to the change of Cl^-^ concentration in blood from the addition of CaCl_2_, as these concentrations affect the electroactive window^80,81^. Considerable peak reduction was observed for the PEG-coated probe (50.8 ± 16%), whereas the F50-C50 hydrogel coating preserved current intensity (76.6 ± 21%) in these assays (**Fig. 4g**). Uncoated, bare devices were also evaluated as controls (44.3 ± 9%) and did not significantly differ from PEG-coated probes (paired t-test, p = 0.931) (**Supplementary Fig. 11, 12**).

Given that our F50-C50 copolymer hydrogel clearly outperformed PEG *in vitro*, we proceeded to test the performance of these two materials *in vivo*. We first sought to evaluate the fouling of sensing probes implanted into the femoral vein of a rat for 5 days (**Supplementary Fig. 13a**). These device bundles comprised two WEs (one PEG-coated and one F50-C50-coated) and one RE (**Supplementary Fig. 13b, c**), and exhibited a diameter of ~500 μm after the hydrogel coating. CV was performed in a (FeCN_6_)^4-^-spiked blood sample *in vitro* to determine the anodic current intensity before and after *in vivo* incubation (a fresh Pt probe was included in the measurements for comparison). After 5 days, the sensor was recovered from the femoral vein (**Supplementary Fig. 13d, e**) and the hydrogel coatings visibly remained intact on the electrode surface (**Supplementary Fig. 13f**). We observed significant signal degradation from bare (81 ± 27% reduction) or PEG-coated (83 ± 23% reduction) probes at the end-point of the study (**Supplementary Fig. 14-15**). Indeed, two thirds of these probes were completely non-functional after five days implanted *in vivo*. In contrast, we could obtain clear signals with notably less signal degradation (24 ± 36% reduction) from all probes coated with our F50-C50 hydrogel (**Supplementary Fig. 14-15**). The results of this study indicated that F50-C50-coated probes performed much more reliably than bare and PEG-coated probes.

### Hydrogel Coating Enables Real-Time Measurement *In Vivo* via DNA Aptamer Biosensors

Based on the *in vivo* pilot study described above, we evaluated these coatings on real-time biosensors that utilize structure-switching aptamers to continuously measure specific analytes *in vivo*^82–85^. Briefly, we engineer the aptamers to undergo reversible structure-switching upon target binding, and we conjugate electroactive reporters (*e.g*., methylene blue, MB) at the distal end of the aptamer (**Fig. 5a**). We continuously measure the electron transfer between the reporter and the electrode using CV, which enables us to monitor the concentration of the analyte in real-time in a reagentless manner in real-time. We and other groups have used such sensors to continuously measure the systemic concentrations of small-molecule drugs *in vivo*^85,88^ and even control their levels using closed-loop feedback control^87^, but biofouling of the surfaces of these devices limits their signal quality, and ultimately, their lifetime^88^. We therefore sought to test whether our top-performing hydrogel coating could improve the performance of aptamer-functionalized electrodes *in vivo*.

**Figure 5.**
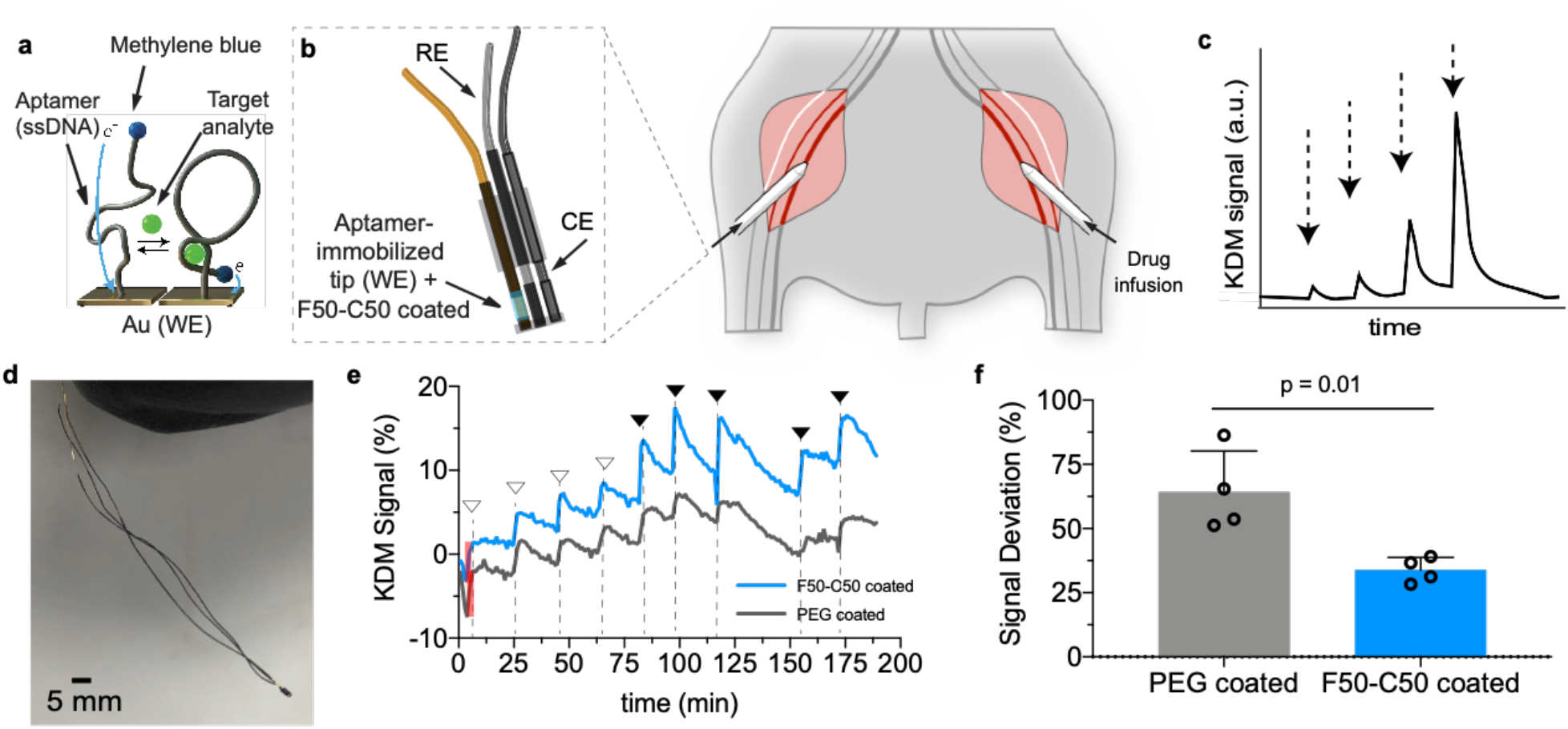
F50-C50 hydrogel coating of a DNA aptamer biosensor enables continuous real-time monitoring of kanamycin *in vivo*. **a.** DNA aptamers that bind kanamycin with high specificity were functionalized onto the tip of Au probes. **b.** The probe was inserted into the femoral vein of a rat through a catheter. Kanamycin was infused into the other femoral vein. **c.** The kanamycin signal response should exhibit fast kinetics and dose-dependence. **d.** Image of the aptamer probe device**. e.** Representative graph of real-time *in vivo* sensing of kanamycin after administration of varying concentrations of the drug by injection into the contralateral femoral vein. Filled arrows represent a 200 μL injection of 200 mM kanamycin and empty arrows indicate a 100 μL injection of 200 mM kanamycin. Shaded red box represents peak intensity of initial baseline signal. **f.** Signal intensities as indicated by peak heights were measured and normalized to baseline signal of each probe. Deviation from the baseline signal between PEG- and F50-C50-coated sensors showed the ability of the F50-C50 hydrogel to maintain higher signal. Data represents mean ±standard deviation, n = 4, unpaired t-test.

As a model system we employed aminoglycoside aptamers that bind kanamycin, which is a widely used small-molecule antibiotic drug (MW = 484.5 Da; diameter ~ 1 nm)^89^. To reduce the diffusion time of the drug to the probe surface, we reduced the thickness of the hydrogel coating by dip-coating the electrodes with precursor solutions for F50-C50 and PEG hydrogels and immediately cross-linked by photopolymerization (**Supplementary Fig. 16**). The mesh size of the resulting hydrogel coating was determined to be 2.33 ± 0.07 nm using FRAP spectroscopy, thus allowing free transport of kanamycin. We performed three experiments with this system. We initially characterized platelet adhesion on probes following incubation in Na_2_EDTA-treated human whole blood for three days at room temperature. We observed significantly less platelet adhesion on F50-C50-coated probes than PEG-coated or bare probes (**Supplementary Fig. 17**). Next, aptamer probes were inserted into a circulating blood flowing system for 6 and 12 days (**Supplementary Fig. 18a**). At the end of each assay, the probes were analyzed by SEM, which showed that the F50-C50 hydrogel coatings had noticeably less platelet adhesion after both timepoints than control probes (**Supplementary Fig. 18b**).

Finally, to test the functional efficacy of these hydrogel coatings *in vivo*, we implanted the aptamer device into the femoral vein of a rat as described above (**Fig. 5b**). Kanamycin (0.1 mL or 0.2 mL at 200 mM) was administered intravenously into the contralateral femoral vein at different timepoints. Aptamer activity was measured using square-wave voltammetry (SWV) at 400 and 60 Hz, and we used the kinetic differential measurement technique to compensate for drift as described previously^86^ (**Fig. 5c, d**). After establishing equilibrium over the course of 60-120 minutes after implantation, we successfully recorded the real-time drug concentration *in vivo* for the first time with F50-C50-coated devices (**Fig. 5e, Supplementary Fig. 19**). Signal peaks in response to kanamycin administration were determined at various time points over the course of roughly 200 minutes following incubation, and normalized to initial peak intensity, due to sensor-to-sensor variations. In these assays, PEG-coated sensors had an average deviation from original signal of 64.2 ± 16.1%, while the enhanced anti-biofouling capabilities of F50-C50 coatings resulted in significantly less deviation, 33.8 ± 4.9%, from the original signal (**Fig. 5f**) (mean ± standard deviation; n = 4, data was analyzed via unpaired t-test where p = 0.01).

## DISCUSSION

In this work we developed a library of 172 polyacrylamide copolymer hydrogels, the most extensive library of this type generated to date, to discover anti-biofouling coatings for biomedical devices that outperform current gold standard materials. Machine-learning analysis of high-throughput, parallel platelet fouling assays suggested that the relevant mechanisms underlying the observed anti-biofouling properties of these materials arise from steric and inertial shape properties of the monomers used in these hydrogels. We also demonstrated that the top-performing formulation exhibits mechanical properties resembling those of human arteries. Finally, we validated the utility of this hydrogel formulation for improving the performance of biosensor devices both *in vitro* and *in vivo*.

To the best of our knowledge, only a small subset of polyacrylamide polymers have been explored previously as candidates for anti-biofouling coatings, including those derived from acrylamide^54,90^, hydroxyethylacrylamide^52,91^, (3-methoxypropyl)-acrylamide and N-isopropyl-acrylamide^92^. Polyacrylamide-based polymers are highly stable over long time-frames under physiological conditions, thereby enabling their use with medical devices requiring long functional lifetimes^93,94^.

While PEG and zwitterionic polymers are considered gold standard materials for anti-fouling coatings, our platelet screening assay identified numerous polyacrylamide copolymer hydrogel formulations exhibiting superior anti-biofouling performance, highlighting the promise of polyacrylamide derivatives in these applications.

Our selected formulation for the L100 PEG hydrogel (PEG diacrylate Mn 780 and PEG acrylate 480) was chosen so as to design a PEG hydrogel that did not meaningfully differ in stiffness from the other hydrogels in our library at the same polymer content. As other PEG monomers have also been used historically for anti-biofouling studies, namely PEG diacrylates and PEG dimethyacrylates of various molecular weights, we evaluated three alternative formulations: (i) PEGDMA740 (15wt%) plus PEGMA550 (5wt%), (ii) PEGDA10K (20wt%), and (iii) PEGDMA10K (20wt%). Each of these formulations comprised 20wt% solids and 1wt% photo-initiator. We found that these formulations exhibited shear storage modulus values ranging over two orders of magnitude (**Supplementary Fig. 20**) and resembling values determined for the library of polyacrylamide-based hydrogels (**Fig. 1e**). Further, we found that these alternative PEG hydrogels exhibited higher IgG and fibrinogen adsorption than the primary L100 PEG control (**Supplementary Fig. 21a**), while platelet adhesion was similar for all formulations evaluated (**Supplementary Fig. 21b**). For this reason, we were confident in our use of L100 as a representative PEG formulation in all of our subsequent experiments.

The anti-biofouling behavior of the various copolymer formulations was evaluated using machine learning algorithms with several different feature sets to elucidate the molecular features that give rise to the observed behavior. The features that were common amongst the top performing hydrogel formulations (Feature Set B) were heavily represented by connectivity and shape descriptors. Coupled with other studies^58,65,66^ indicating that ligand mobility makes an entropic contribution to anti-fouling performance, this work provides critical insight into the molecular mechanisms underpinning anti-biofouling behaviors and potentially enabling the rational design of next generation coatings. Recently, Rostam *et al*. developed a (meth)acrylate and (meth)acrylamide polymer library to modulate the foreign body response to implanted materials and made use of machine learning approaches to analyze important chemistry descriptors driving macrophage phenotype and implant fouling. While this study made use of arrays of neat polymers instead of hydrogels, steric (and electronic factors) were also attributed to feature importance^35^.

We sought to eliminate surface topography as a contributor to differences in biological performance in the PAAm library by polymerizing the hydrogels between glass slides. Although SEM images revealed that the various hydrogel formulations evaluated exhibit different levels of activation of platelets (**Supplementary Fig. 3**), the differences in hydrogel surface topography that were apparent in SEM are attributable to the sample processing, wherein hydrogels were lyophilized prior to imaging. To evaluate the true topology of the hydrated hydrogel surfaces, we performed optical microscopy on a subset of materials, including our top-performing polyacrylamide formulation, homopolymer hydrogels that comprise the top formulation, a subset of various polyacrylamide hydrogels across the library, and control materials. Optical microscopy images (**Supplementary Fig. 22a-g**) revealed that all of the materials evaluated are relatively flat, with << 3 μm variation in their height profile, and have negligible surface roughness (**Supplementary Table 5**). As the topography and surface roughness of the hydrogels are similar across all materials evaluated, we believe surface topography is not an important variable within the hydrogel library.

When the top-performing polyacrylamide copolymer hydrogel was used as a coating on an electrochemical biosensor device, we observed little reduction in signal *in vitro* in the presence of whole blood. The mechanism behind this improved signal could be attributed to two critical factors that are enhanced by the hydrogel. First, the surface of a bare electrode is exposed to a broad spectrum of molecules, leading to broad peaks, while the pores of the hydrogel coating exclude molecules too large to diffuse through the matrix, reducing peak widths observed with hydrogel-coated probes. Second, the intensity of measured peaks is maintained throughout the duration of the assay with F50-C50 hydrogel-coated probes, suggesting that this formulation precludes biofouling^59,95^ while allowing molecules of interest to diffuse through the hydrogel matrix and bind reversibly to the surface to produce signal. In contrast, PEG-coated probes lose their ability to sense molecules over time, despite having a comparable mesh size to the F50-C50 hydrogel coating, suggesting that free diffusion of molecules through the coating is impaired by fouling. Indeed, other studies have suggested PEG coatings may not be suitable for use on biosensors of this type as they can hinder electron transfer^96^.

Through in vivo implantation studies, we observed that F50-C50-coated probes performed much more reliably than bare and PEG-coated probes. Indeed, only one probe from each of the control groups showed any response after five days implanted in the femoral vein of a rat on account of fouling in vivo. In contrast, all three F50-C50-coated probes exhibited clear signals with higher anodic currents (Supplementary Fig. 13-15). Similarly, PEG-coated DNA aptamer sensors showed twice as much deviation from their original signal relative to F50-C50-coated sensors within the first two hours after implantation in the femoral vein of a rat (Fig. 5). These studies suggest that the majority of signal loss likely occurs in the first few hours of blood exposure and changes relatively little out to 5 days following implantation. While future work will certainly be needed to evaluate the function of F50-C50-coated sensors over longer durations relevant for certain applications (e.g., catheter-bound sensors), these studies suggest that performance of F50-C50-coated sensors is rather stable through the first 5 days, thereby exhibiting promise for maintaining sensor function over longer time-frames. Overall, these data show that combinatorial screening of copolymer hydrogels offers an exciting avenue for the discovery of anti-biofouling coatings.

Although it may not be intuitively obvious from a materials design point-of view, we demonstrate that certain copolymer formulations can deliver considerably better anti-biofouling performance in comparison to gold standard materials than homopolymers. These observations highlight the value of coupling chemical design intuition with high-throughput screening approaches for discovery of new materials. We have demonstrated the applicability of these new hydrogel coatings to the surface of electrodes using facile synthetic approaches that are amenable to scalable manufacturing, although future studies must be conducted to optimize the adhesion strength of these hydrogels to device surfaces for longer-term biomedical use^97,98^ and to enhance the durability of the hydrogel materials^99^. With further improvements, we believe this strategy of using combinations of acrylamide-based monomers may enable the discovery of anti-biofouling materials enabling long-term implantation of biosensor devices to continuously monitor chronic biomedical conditions.

## METHODS

### Materials and Reagents

All reagents were purchased from Sigma-Aldrich and used as received, unless later specified. Au, Pt, and Ag wires were purchased from A-M Systems. EDTA-treated human whole blood for flowing *in vitro* measurements was purchased from BioIVT. Kanamycin monosulfate was ordered in USP grade from Gold BioTechnology.

### Hydrogel Preparation

Pre-polymer formulations containing 20 wt% acrylamide monomer, 1 wt% lithium phenyl-2,4,6-trimethylbenzoylphosphinate (LAP) as photo-initiator, and 1 wt% methoxy-bisacrylamide were mixed and pipetted between two glass slides separated by a silicone spacer (0.25 mm ± 0.05 mm). Gels were cross-linked in a Luzchem photoreactor system with 8 W bulbs and an intensity of 25-40 W/m^2^ (LZC-4, *hv* = 350 nm, 15 min). Due to swelling of polyacrylamides in water, they were placed in 1x PBS for at least 24 hr before being punched with a 6 mm biopsy punch. PMPC, PSBMA, and PHEMA hydrogels were prepared as described previously^100,101^. Briefly, polyzwitterionic hydrogels were prepared with monomeric solutions in 1 M NaCl with 4% N,N’-methylenebisacrylamide (MBAm) cross-linker. A stock solution of 15% sodium metabisulfite and 40% ammonium persulfate was added to the prepolymer solution at ~1wt% and polymerization was initiated at 60 °C. PHEMA hydrogels were prepared in a solution of ethanol : ethylene glycol : water (1:1.5:1.5) with tetraethyleneglycol dimethacrylate as a cross-linker. A stock solution of 40% ammonium persulfate and 0.5 wt% TEMED was added to the prepolymer solution at 1.5 wt% and polymerization was initiated at 60°C. PEG hydrogels were prepared similarly with a prepolymer solution of 15 wt% PEG diacrylate (M_n_ 780^102–104^) and 5 wt% PEG acrylate (M_n_ 480). Prior work has shown that PEG molecular weight (M_n_ > 10K) affects anti-fouling performance when PEG is presented as brush polymers tethered to a surface. In addition, when molecular weight of PEG exceeds 40 kDa, issues arise such as potential immunogenicity and accumulation in tissues^105^. Here, we have used hydrogels to create a fully cross-linked network and, to our knowledge, have not seen effects of PEG molecular weight.

Additional PEG hydrogels were synthesized with 20 wt% PEG10K diacrylate and 1 wt% LAP (PEGDA10K), 20 wt% PEG10K dimethacrylate and 1 wt% LAP (PEGDMA10K) and 15 wt% PEG740 dimethacrylate, 5 wt% PEG 550 methacrylate and 1 wt% LAP (PEGMA740_PEGMA550). Prepolymer solutions were mixed and pipetted between two glass slides separated by a silicone spacer (0.25 mm ± 0.05 mm). Gels were cross-linked in a Luzchem photoreactor system with 8 W bulbs and an intensity of 25-40 W/m^2^ (LZC-4, *hv* = 350 nm, 15 min). Hydrogels were placed in 1x PBS for at least 24 hr before being punched with a 6 mm biopsy punch.

### Hydrogel Synthesis Modifications

All 2-acrylamido-2-methyl-propane sulfonic acid (AMPSAm) formulations were made with slightly basic solution of 2:3 1 M NaOH:water. [tris(hydroxymethyl)methyl]acrylamide (tHMAm) formulations were made with 50:50 dimethylformamide:water as well as 100% N-isopropylacrylamide (NiPAm), 75% diethylacrylamide (DEAm)/25% NiPAm, and 25% hydroxymethylacrylamide (HMAm)/75% NiPAm. Acrylamidopropyl)trimethyl-ammonium (APTAC)-based formulations were used with 2-3x MBAm concentration to achieve comparable mechanical properties.

### Platelet Adhesion Test

Fresh whole rat blood was mixed in a 10:1 volume ratio of an acid citrate dextrose (ACD) anticoagulant buffer (containing 2.13% free citrate ion, BD Vacutainer Specialty Venous Blood Collection Tubes) for the preparation of platelet-rich plasma (PRP). PRP was obtained via centrifugation at 600 x g for 10 min at 10 °C. The platelets were counted using a Countess II Automated Cell Counter (ThermoFisher Scientific) and diluted to 2.5 x 10^6^ platelets/mL in 1x PBS. 6 mm punches of hydrogels were placed in ultra-low adhesion 96-well plates and incubated for 24 hr at 37 °C. Gels were UV sterilized for 5 min prior to incubation with platelets. 100 μL of PRP was pipetted on top of the hydrogels. The plates were placed on a rotary shaker for 1 hr at room temperature. Platelets were rinsed once with 1x PBS and fixed with 4% paraformaldehyde (PFA). Cells were imaged with an EVOS XL Core Imaging System microscope (Life Technologies).

### Platelet Detection in Images

Platelets in the images are small round objects ~3-4 μm in diameter. Platelet images are of different color and can include noise such as gel chunks or dust. The noise typically exhibits sizes considerably smaller or larger than platelets. The platelets in the images were detected using a difference-of-Gaussian approach by blurring images using Gaussian kernels of a range of standard deviations in increasing order. Stacks of the differential images between two successively blurred images form a cube, and where blobs are local maxima of intensity. Blobs of noisy objects are avoided by tuning the range of standard deviations used in the process.

### IgG and Fibrinogen Adhesion

Hydrogel discs (n = 3 for each sample) were punched into a low-adhesion clear 96-well plate. To measure IgG adsorption to the surface of the hydrogels, 100 μL of 2 μg/mL Alexa Fluor 488-AffinityPure Rabbit Anti-Bovine IgG (Jackson ImmunoResearch) was added to each well; the plate was then sealed and placed on a shaker plate overnight. For fibrinogen adhesion, 100 μL of 5 μg/mL fibrinogen from human plasma conjugated to AF488 (Invitrogen) was added to each well; the plate was then sealed and placed on a shaker plate for 1 hour. Samples were then washed 5x with PBS, after which the gels were transferred to a black 96-well plate and fluorescence readings were taken (BioTek Synergy H1 microplate reader).

### Fluorescence Recovery After Photobleaching (FRAP)

Hydrogel samples were loaded with 0.5 wt% FITC-dextran (4kDa). An Inverted Zeiss LSM 780 laser scanning confocal microscope (Germany) with a Plan-Apochromat 20X/0.8 M27 objective lens was used for FRAP analysis. We used pixel dwell time 1.58 μs. Samples were photobleached with 405-nm and 488-nm argon lasers set to 100% intensity. The samples were placed in a sterile 0.16-0.19 mm thick glass bottom μ-dish (MatTek). ZEN lite (Zeiss) software was used for all FRAP tests. To avoid any extra noise, the high voltage was limited to 700 V. Different tests (n = 5) were made at different locations of the sample. For each test, 10 control pre-bleaching images were captured per second, and we then bleached the spot with a pixel dwell time of 177.32 μs. 390 post-bleaching frames were recorded per second to form the recovery exponential curve. The pixel size was set to be 0.83 μm. The diffusion coefficient was calculated as described in the literature^106^: D= **γ**_D_(**ω**^2^/4**τ**_1/2_) where **γ**_D_ = 0.88 for circular beams, **ω** is the radius of the bleached ROI (12.5 μm), and **τ**_1/2_ is the half-time of the recovery. To estimate the mesh size (**ξ**) of our hydrogels, we used the obstruction theory of Amsden *et al*.^107^:

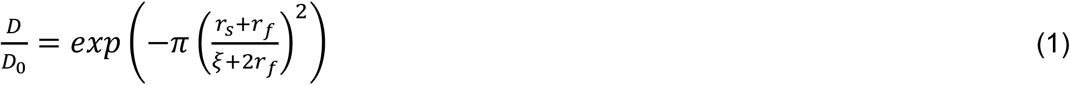

where *D* is diffusivity of the solute in the hydrogel, *D*_0_ is the diffusivity of the solute within the liquid in the hydrogel (saline-sodium citrate buffer), which is assumed to be the same as in pure water, *r_s_* is the radius of the solute (3.51 nm for FITC-dextran 4kDa), *r_f_* is the radius of the polymer chains within the hydrogel network (estimated as 0.65 nm^108^ for F50-C50 hydrogel, 0.51 nm for PEG^109^, 2.00 nm for pHEMA^110^, 1.00 nm for PMPC^111,112^ and 0.9 nm for PCBMA^113^). The diffusivity (*D*_0_) of a solute in a pure liquid was determined with the Stokes-Einstein equation to be 69.91 μm^2^/s using a solution viscosity of **η** = 0.89 x10^-4^ Pa s.

### Scanning Electron Microscopy

SEM images were acquired with an FEI Magellan 400 XHR microscope with a beam voltage of 1 kV. The sample was lyophilized prior to imaging, pressed onto silver paint and sputter-coated with Au:Pd (60:40) before imaging. When imaging hydrogels and adherent platelets, samples were fixed with 4% glutaraldehyde (GFA) for 5 minutes, then dehydrated with 10%, 30%, 50%, 70%, 90%, 100% ethanol solutions for 10 minutes each. Samples were then lyophilized, gently pressed onto silver paint and sputter-coated before imaging.

### Performance Classification

The relevant performance range of the antifouling polymer is platelet count under 100 units. This requirement naturally sets up the data analytic task as classification. From this point, antifouling candidates are considered as members of one of two performance classes. The positive class includes polymers with median platelet count below or equal to 100, and the negative class includes polymers with median platelet above 100 counts. Out of the characterized polymers, there were 76 members in the positive class and 91 members in the negative class. The slight class imbalance was considered irrelevant for practical purposes, and can be mitigated using standard techniques such as over- and under-sampling. The data was organized into three different feature sets (A, B, C). Feature Set A was reflective of the molar ratios of the monomers in the respective polymer. Feature Set B increased granularity by generating vectors of physical-chemical descriptors to the monomers and weighting these by the molar ratios of the monomer in the polymers. A simplified version was generated in Feature Set C, by summing up descriptor vectors of the monomers weighted by their molar ratios in the corresponding polymers. These classifiers (**Supplementary Table 3**) showed highest performance with Feature Sets A and B.

### Machine-Learning Data Analysis

Classification was performed using a random forest (RF) model, as implemented in *scikit*-learn 0.20.2. As a model, RF offers robustness to overfitting and the capability to evaluate relevance of data features. RF model parameters were selected via cross-validated grid-search; optimized parameters included the number of estimators, maximum estimator depth, and maximum number of features used in the splitting procedure. Three-fold cross-validation was performed over the training set, which included 75% of the available observations. The model produced as a result of the cross-validated grid search was tested on the withheld test set that included 25% of the available observations. We selected MDA to evaluate feature relevance, which captures the drop in accuracy of the trained model under random permutation of the feature whose relevance is being evaluated with the rest of the features. Higher positive values of MDA are indicative of a larger drop in accuracy in the permutation test and point to higher impact of the respective feature on the model predictions. MDA values close to zero are indicative of minimal impact of the feature permutation on accuracy, and help identify irrelevant features. Negative MDA values point at features whose permutation increases model accuracy. Such features do not contribute meaningfully to the model performance. Model performance on the test set was evaluated using standard measures of accuracy in classification tasks, such as recall, accuracy, and f1-score.

### Expanded Analysis of Platelet Homogeneity

The coverage profile for each formulation was generated in the following manner: (i) partition the image of the polymer surface into square bins, (ii) count the platelet population in each bin, (iii) generate a population histogram for all images obtained for a given formulation, and (iv) normalize the histogram to the total number of population bins obtained for a given formulation. There are several general types of population histograms that one might expect to encounter (**Supplementary Fig. 4a–d**). We recognize that the difference between homogenous and heterogenous coverage modes is quantitative, not qualitative. In both cases, histograms correspond to distributions that have tails; the thickness of the tails distinguishes the two cases, where the tails fall off faster in homogenous cases than in heterogenous cases.

Using normalized population histograms as features, we performed anomaly detection analysis with the Isolation Forest algorithm^114^. Each histogram received an anomaly score; negative values indicate outliers, and positive values indicate inliers. The notions of inliers and outliers are reflective of the structure of the dataset and do not have absolute meaning. The anomaly scores are visualized by reducing dimensionality of the dataset to two dimensions via t-SNE manifold embedding and encoding the anomaly score of the samples as the color (**Supplementary Fig. 4i**). Outlier analysis identifies histograms corresponding to homogenous coverage as the most normal. The change of the anomaly score towards the most abnormal system is associated with increased heterogeneity of the coverage (**Supplementary Fig. 4j**). Overall, there were 121 systems that bore signs of increasing heterogeneity of coverage, suggesting that the steric mode of fouling plays a meaningful role.

### Rheological Characterization

Oscillatory rheology measurements were performed with a TA Instruments AR-G2 rheometer. Amplitude sweeps were conducted at a frequency of 10 rad/s from 0.1-100%. Frequency sweeps were conducted at 0.1% strain from 0.1-100 rad/s. All tests were performed at 25 °C using an 8 mm parallel plate geometry.

### Tensile Test

Tensile strength measurements were performed with an Instron series 5560A with 100 N load cell. Tensile tests were conducted at 0.2 mm/s at room temperature.

### Nanoindentation Test

Young’s modulus measurements of the hydrogels were performed using a Piuma Nanoindenter (Optics11) with a probe from the same manufacturer with a stiffness of 38.8 N/m and a tip radius of 27.0 μm. Calibration was conducted as on glass under wet conditions as per the manufacturer’s instructions. Each sample was immersed in a sterile saline buffer solution (Opti-free - Replenish; Alcon) before the nanoindentation experiments. The indentation depth was fixed to be <1 μm in order to avoid bottom effects. At least 8 force curves were used to determine the local Young’s modulus of each sample, using the Optics11 Nanoindenter V2.0.27 software. The results are shown as mean ± standard error of the mean. A Hertzian contact model parameter was used for the fit of the curves assuming that the Poisson’s ratio of the samples is **υ** = 0.5.

### Fabrication of Electrochemical Biosensors for *In Vitro* Assessment

The working electrode (WE) was prepared with 75-μm diameter gold wire (PFA-coated). Ag/Ag Cl wires were prepared by incubating silver electrodes in bleach solution, rinsed vigorously with water, and dried. Pt wire was bundled with the Au electrode after bundling with Ag/AgCl wire to prevent shorting. The tip of the device was cut with a razor blade to expose bare gold and rebleached. No surface roughening was applied. To prevent wires from touching the side of the Eppendorf tube walls, they were propped in the middle of the tube. A hole was drilled in the cap of a 0.3 mL Eppendorf tube to fit 0.04” Tygon tubing. One inch of tubing was inserted into the drilled hole, and we applied epoxy and cured with UV light (365 nm) for 20 seconds. Wires were then inserted through the Tygon tubing into the tube.

### Hydrogel Coatings on Electrochemical Biosensors for *In Vitro* Assays

1 μL of prepolymer solution was pipetted into a 100 μL pipette tip. The probes were inserted and polymerized for 5 seconds was photo-cured at 365 nm. The pipette tip was removed, and the hydrogel coating was visible by eye. The hydrogel was then photo-cured for an additional 30 seconds.

### *In vitro* Assays with Electrochemical Devices

Bare, PEG-coated, and F50-C50-coated probes were incubated in EDTA-treated human whole blood at room temperature with 250 μL of 15 mM (FeCN_6_)^4-^-to record baseline CV data. This step was limited to 5 minutes to ensure a steady-state readout while minimizing biofouling. Probes were rinsed in 1x SSC buffer for 5 minutes to thoroughly remove any intrinsic fouling and transferred to 250 μL whole blood, followed by addition of CaCl_2_ to a final concentration of 50 mM. After 2 hr, probes were transferred to the original tube used for baseline measurement. The device was incubated for 5 minutes to allow the diffusion of (FeCN_6_)^4-^ throughout the hydrogel. CV scanning was performed on the Au wire (potential range: −0.7–0.8 V, step size: 1 mV, scan rate: 0.1 V/sec).

### Fabrication of Electrochemical Biosensors for *In Vivo* Assessment

Bare gold was cut into ~10-cm wires. The tip (~1 mm) was bundled with black medical heat shrink tubing twice (ID 0.006” ± 0.001, Nordson Medical 103-0325). Another two layers of heat shrink were applied ~1-2 mm away, allowing exposure of ~1–2 mm of bare gold. Ag/AgCl wires were bundled likewise to prevent shorting. Two gold wires (WE) and one Ag/AgCl wire (RE) were bundled at the tip with clear shrink wrap (ID 0.02” ± 0.001, Nordson Medical 103-0249) and at the middle of the probe to provide mechanical strength when pushing through the catheter.

### Hydrogel Coatings on Electrochemical Biosensors for *In Vivo* Assays

For each device, one Au wire was left bare and one was coated with hydrogel. The prepolymer solution of hydrogel was pipetted into a short segment of tubing from a 26G catheter with inner diameter of 0.6 mm. The device was pushed through the tubing and aligned with one of the two bare gold wires. UV light (365 nm) was applied for 5 seconds and the tubing was gently removed. The hydrogel coating was visible on the entirety of one Au probe only, and UV light was applied for 25 more seconds to ensure complete photopolymerization. Probes were stored in SSC buffer and placed in 200 μL of EDTA-treated rat blood (BioIVT) with 15 mM (FeCN_6_)^4-^ to record baseline data. Probes were incubated for 5 minutes to allow diffusion of (FeCN_6_)^4-^. CV scanning was performed on the Au wire (potential range: −0.7–0.8 V, step size: 1 mV, scan rate: 0.1 V/sec).

### *In Vivo* Assays with Electrochemical Devices

Live animal studies were performed with male Sprague–Dawley rats under Stanford APLAC protocol number 33226. All rats used in this work were purchased from Charles River Laboratories at a weight of 350-475 g. The rats were anesthetized using isoflurane gas (2.5%) and monitored continuously. After exposing the femoral vein, a 24G catheter was implanted into the vein for sensor probe insertion. The needle of the catheter was removed, and the catheter was clipped. The devices were inserted through the catheter and pushed far into the vein. The catheter was clamped, and the device was secured in place, after which the incision was sutured. At the end of the experiments, animals were anesthetized using isoflurane gas and the devices were removed for immediate measurement in blood used for baseline sample. Rats were then euthanized by exsanguination while under general anesthesia.

### Aptamer Device Fabrication and Functionalization

The kanamycin aptamer probes were synthesized by Biosearch Technologies with the following sequence: 5′-HS-(CH2)_6_-GGGACTTGGTTTAGGTAATGAGTCCC-MB-3′. Probes were thiolated at the 5′ end with a 6-carbon linker for self-assembly onto the WE, and conjugated with a methylene blue (MB) redox label at the 3′ end with a 7-carbon linker for charge transfer measurements. The modified DNAs were purified through dual HPLC by the supplier. Upon receipt, the construct was dissolved and diluted to 100 μM using UltraPure water (Thermo Fisher Scientific) and frozen at −20 °C in individual aliquots of 1 μL until use. The WE was prepared with 8 cm pure Au wire (75 μm diameter) insulated with heat-shrinkable tubing (Nordson Medical, 103-0325) to define the aptamer immobilization surface. The exposed gold wire has a length of 1~2 mm with an overall surface area of 0.25~0.5 mm^2^. No surface roughening was applied. Before immobilizing the aptamer probes, the wire was rinsed with acetone, ethanol, and deionized water in a sonicator sequentially, followed by electrochemical cleaning. CV scanning on the gold wire was performed in 500 and 50 mM sulfuric acid solutions (potential range: −0.4–1.5 V, step size: 1 mV, scan rate: 0.1 V/sec), each with three scans. An aliquot of the DNA construct was thawed and reduced for 40 minutes with 2 μL 100 mM tris(2-carboxyethyl)phosphine at room temperature in the dark. The reduced DNA construct was diluted to 1 μM with deionized water, and a freshly cleaned gold electrode was immersed for 1 h at room temperature in dark. Next, the sensor was rinsed with deionized water for 3 min, followed by immersion in 6 mM 6-mercapto-1-hexanol in 1x SSC buffer for 2 hr at room temperature in the dark to passivate the remaining gold surface and remove nonspecifically adsorbed DNA. The sensor was rinsed with deionized water for another 3 min and stored in 1x SSC buffer at 4°C for 12 h before application of the hydrogel.

Electrochemical measurements were conducted using an EmStat Blue instrument (Palm Instruments BV) in square-wave voltammetry (SWV) mode. As only working and reference electrodes are employed in the device, the input connections for the reference and the counter electrode from the electrochemical analyzer were shorted. The WE was scanned in continuous succession with a scan period of 2 seconds, alternating between two SWV frequencies (400 and 60 Hz) at a constant SWV amplitude of 36 mV. Two frequencies were used in order to apply KDM for drift mitigation. As the MB redox peak was typically observed at about −350 mV in our setup, a potential range of −500 to −100 mV (with respect to the Ag/AgCl reference) was selected during the SWV scan. A custom peak-fitting script was created to fit the SWV measurements with a Gaussian curve on a hyperbolic baseline. Peak currents were then normalized to a baseline peak current to generate the signal gain. All reported gains were obtained via KDM, with the difference divided by the average of gains from the 400 and 60 Hz signals.

### Hydrogel Coatings on Aptamer Devices

Acrylamide monomers were purified through a basic alumina column. The hydrogel was applied to the sensing gold wire through capillary force, in which the sensor was dipped into the hydrogel prepolymer solution for 5 sec and removed immediately. The hydrogel was then photopolymerized by applying 365-nm UV light for 30 sec. No reduction in the MB peak current was observed after UV application. An Ag/AgCl reference electrode (75-μm-diameter Ag wire chlorinated in bleach overnight) was attached to the hydrogel-coated gold wire using heat shrinkable tubing. The final device was placed in 1x SSC buffer at 4 °C overnight before use.

### *In vitro* Assessments of Aptamer Devices in Stationary and Flowing Blood

Aptamer devices were incubated in EDTA-treated human whole blood (Bio-IVT) or inserted via a catheter into a closed-loop flowing blood set-up (Masterflex L/S Easy-Load II EW-77200-80 pump head at 1.5 mL/min) for 6 or 12 days at room temperature (**Supplementary Fig. 18**). Probes were then immersed in 2.5% GFA for 15 minutes and rinsed with PBS. Probes were then freeze-dried, lyophilized, and coated with Au:Pd for SEM imaging.

### *In vivo* Assessment of Aptamer Devices

Live animal studies were performed using Sprague–Dawley rats under Stanford APLAC protocol number 33226. Male rats were purchased from Charles River Laboratories at a weight of 300 - 350 g. The rats were anesthetized using isoflurane gas inhalation (2.5%) and monitored continuously. After exposing both femoral veins, a 20G catheter was implanted into the left femoral vein for sensor probe insertion, whereas a 22G catheter was implanted into the right femoral vein for drug infusion. 0.1 - 0.3 mL of heparin (1000 U/mL, SAGENT Pharmaceuticals) were immediately infused through the catheter to prevent blood clotting. The sensor was secured in place with a surgical suture after the insertion and allowed to equilibrate for 1-2 hours. A sequence of bolus injection was performed manually using a 5 mL syringe. In each administration, 100 or 200 μL of 200 mM kanamycin in PBS buffer was injected through the sensor-free catheter. At the end of the experiments, animals were euthanized by exsanguination while under general anesthesia.

### Degradation Assay for Hydrogels

Solutions of 12 M HCl, 12 M NaOH, 30% peroxide, and 1 mg/mL Amano Lipase PS were prepared (Sigma-Aldrich). Hydrogel discs were placed into one of each of these solutions at 50 °C for up to 72 hours. At specified time intervals, swollen hydrogel discs were removed from the solution and placed into deionized water for 24 hours to remove salts. Samples were then freeze-dried and lyophilized before mass was recorded. Weights were normalized to mass at time 0 to determine mass degradation.

### Optical Microscopy and Roughness of Hydrogels

Differential interference contrast (DIC) miscroscopy (10X) of the gel surfaces was performed with a Keyence VK-X250K/260K 3D laser scanning confocal microscope. We used a 408-nm violet laser and performed image analysis with VK Viewer and MultiFileAnalyzer software from Keyence. Surface roughness measurements used to quantify surface roughness measurements were defined and parameterized by ISO 25178 and measuring height profiles across the x-y plane of the hydrogel samples.

### Statistical Analysis

All values of significance were determined using a one-way ANOVA or student’s t-test with Prism GraphPad 8.4 software. As we noted drastic differences in clotting time between batches of blood, a paired t-test was used to determine significance between PEG-coated and F50-C50-coated probes in **Fig. 4**. In **Fig. 5**, an unpaired t-test was performed to compare performance of PEG-coated and F50-C50-coated sensors as there was one only group per leg of each animal (no pairing). In **Supplementary Fig. 15** and **Supplementary Fig. 21**, a one-way ANOVA test was used as there were multiple groups being compared.

## Ethical Statement

All animal procedures were performed according to Stanford APLAC approved protocols.

## Data Availability

All data supporting the results in this study are available within the article and its Supplementary Information. The broad range of raw datasets acquired and analyzed (or any subsets thereof), which would require contextual metadata for reuse, are available from the corresponding author upon reasonable request.

## Code Availability

All code that supports the findings from this study is available on reasonable request.

## Acknowledgements

This work was supported in part by the NIDDK (R01DK119254). Doreen Chan is grateful for an award by the Department of Defense, Air Force Office of Scientific Research, National Defense Science and Engineering Graduate (ND-SEG) Fellowship, 32 CFR 168a with government support under FA9550-11-C-0028. Dr. Eneko Axpe is thankful for funding support from a Stanford Bio-X Interdisciplinary Initiative Program Round 8 (2016) Seed Grant and the European Commission for the Marie Curie global fellowship. Part of this work was performed at the Stanford Nano Shared Facilities (SNSF), supported by the National Science Foundation under award ECCS-1542152. We thank Grant Shao, Lab Operations Engineer at Stanford Nano Shared Facilities, for invaluable assistance in obtaining optical microscopy images and roughness measurements.

## Author contributions

D. C., J.-C.C., E.A., S.S., V.A.P., D.Yu.Z., H.T.S., and E.A.A. designed experiments. D.C., J.-C.C., E. A., L.B., S.S., V.A.P., D.Yu.Z., C. L. M., A. K. G., J. L. M., H.T.S., and E.A.A. conducted experiments. S.W.B. assisted with surgeries. D.C., J.-C.C., E.A., L.B., C. L. M., H.T.S., and E.A.A. analyzed data. D.C., E.A., S.S., V.A.P., D.Yu.Z., J.-C.C., H.T.S., and E.A.A. wrote the paper.

