## Supplementary Information for "Combinatorial Polyacrylamide Hydrogels for Preventing Biofouling on Implantable Biosensors"

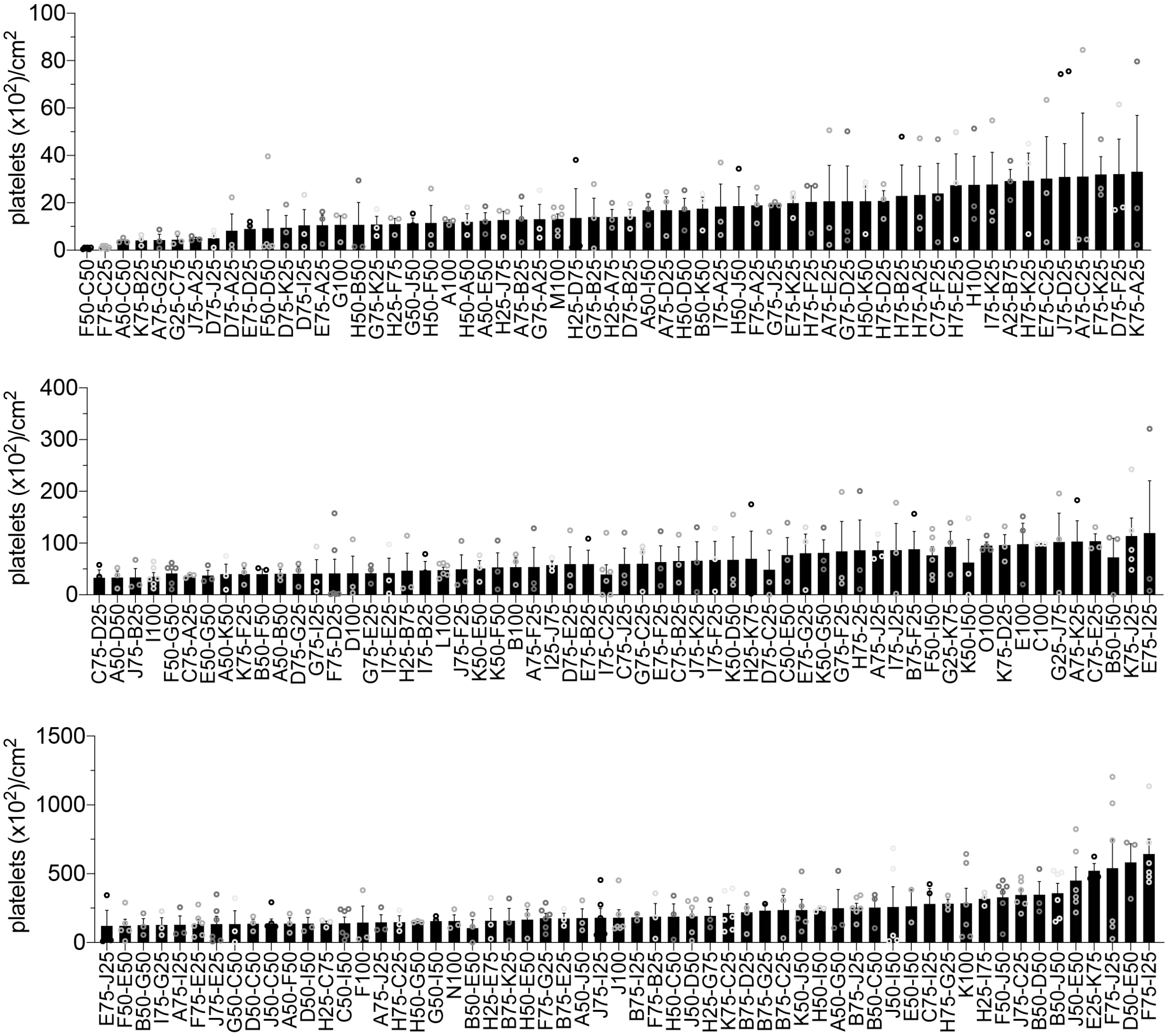

**Supplementary Fig. 1. Platelet adhesion screen.** Combinatorial polyacrylamide hydrogels were evaluated in a high-throughput parallel screen for platelet adhesion. Mean ± standard error platelet counts on hydrogel surfaces are shown with $n \geq 3$.

**Supplementary Table 1.** Abbreviations of acrylamide monomers.

| | Abbreviation | Monomer |
|---|---|---|
| A | Am | acrylamide |
| B | DMAm | dimethylacrylamide |
| C | DEAm | diethylacrylamide |
| D | MPAm | (3-methoxypropyl)acrylamide |
| E | HMAm | hydroxymethylacrylamide |
| F | HEAm | hydroxyethylacrylamide |
| G | tHMAm | [tris(hydroxymethyl)methyl]acrylamide |
| H | ALMP | acryloylmorpholine |
| I | APTAC | (acrylamidopropyl)trimethyl-ammonium |
| J | AMPSAm | 2-acrylamido-2-methyl-propane sulfonic acid |
| K | NiPAm | N-isopropylacrylamide |

**Supplementary Table 2.** List of polyacrylamide formulations in the hydrogel library. Hydrogel formulations generally consist of 20 wt% total monomer, 1 wt% cross-linker, and 1 wt% photo-initiator.

| Ranking from Fouling Assay | Formulation Code | Monomer 1 | Monomer 1 Content (wt%) | Monomer 2 | Monomer 2 Content (wt%) |
|---|---|---|---|---|---|
| 1 | F50-C50 | HEAm | 10 | DEAm | 10 |
| 2 | F75-C25 | HEAm | 15 | DEAm | 5 |
| 3 | H50-B50 | ALMP | 10 | DMAm | 10 |
| 4 | H25-D75 | ALMP | 5 | MPAm | 15 |
| 5 | F50-D50 | HEAm | 10 | MPAm | 10 |
| 6 | D75-A25 | MPAm | 15 | Am | 5 |
| 7 | F75-D25 | HEAm | 15 | MPAm | 5 |
| 8 | A50-C50 | Am | 10 | DEAm | 10 |
| 9 | G25-C75 | tHMAm | 5 | DEAm | 15 |
| 10 | A75-G25 | Am | 15 | tHMAm | 5 |
| 11 | J75-A25 | AMPSAm | 15 | Am | 5 |
| 12 | A75-C25 | Am | 15 | DEAm | 5 |
| 13 | K75-B25 | NiPAm | 15 | DMAm | 5 |
| 14 | D75-J25 | MPAm | 15 | AMPSAm | 5 |
| 15 | H50-F50 | ALMP | 10 | HEAm | 10 |
| 16 | D75-I25 | MPAm | 15 | APTAC | 5 |
| 17 | D75-K25 | MPAm | 15 | NiPAm | 5 |
| 18 | G75-D25 | tHMAm | 15 | MPAm | 5 |
| 19 | G75-A25 | tHMAm | 15 | Am | 5 |
| 20 | A75-E25 | Am | 15 | HMAm | 5 |
| 21 | G75-K25 | tHMAm | 15 | NiPAm | 5 |
| 22 | E75-D25 | HMAm | 15 | MPAm | 5 |
| 23 | G50-J50 | tHMAm | 10 | AMPSAm | 10 |
| 24 | H50-A50 | ALMP | 10 | Am | 10 |
| 25 | I75-A25 | APTAC | 15 | Am | 5 |
| 26 | A50-E50 | Am | 10 | HMAm | 10 |
| 27 | A100 | Am | 20 | - | 0 |
| 28 | J75-D25 | AMPSAm | 15 | MPAm | 5 |
| 29 | H25-A75 | ALMP | 5 | Am | 15 |
| 30 | A75-B25 | Am | 15 | DMAm | 5 |
| 31 | E75-A25 | HMAm | 15 | Am | 5 |
| 32 | H25-F75 | ALMP | 5 | HEAm | 15 |
| 33 | G75-B25 | tHMAm | 15 | DMAm | 5 |
| 34 | H75-A25 | ALMP | 15 | Am | 5 |

| | | | | | |
|---|---|---|---|---|---|
| 35 | D75-B25 | MPAm | 15 | DMAm | 5 |
| 36 | G100 | tHMAm | 20 | - | 0 |
| 37 | H50-J50 | ALMP | 10 | AMPSAm | 10 |
| 38 | H25-B75 | ALMP | 5 | DMAm | 15 |
| 39 | D100 | MPAm | 20 | - | 0 |
| 40 | I75-K25 | APTAC | 15 | NiPAm | 5 |
| 41 | H25-J75 | ALMP | 5 | AMPSAm | 15 |
| 42 | E75-J25 | HMAm | 15 | AMPSAm | 5 |
| 43 | H75-B25 | ALMP | 15 | DMAm | 5 |
| 44 | H50-D50 | ALMP | 10 | MPAm | 10 |
| 45 | A50-I50 | Am | 10 | APTAC | 10 |
| 46 | K75-A25 | NiPAm | 15 | Am | 5 |
| 47 | D75-F25 | MPAm | 15 | HEAm | 5 |
| 48 | H100 | ALMP | 20 | - | 0 |
| 49 | G75-J25 | tHMAm | 15 | AMPSAm | 5 |
| 50 | F75-A25 | HEAm | 15 | Am | 5 |
| 51 | J75-B25 | AMPSAm | 15 | DMAm | 5 |
| 52 | A75-D25 | Am | 15 | MPAm | 5 |
| 53 | B50-K50 | DMAm | 10 | NiPAm | 10 |
| 54 | H75-D25 | ALMP | 15 | MPAm | 5 |
| 55 | C75-F25 | DEAm | 15 | HEAm | 5 |
| 56 | E75-K25 | HMAm | 15 | NiPAm | 5 |
| 57 | A75-F25 | Am | 15 | HEAm | 5 |
| 58 | E75-C25 | HMAm | 15 | DEAm | 5 |
| 59 | G75-I25 | tHMAm | 15 | APTAC | 5 |
| 60 | J75-F25 | AMPSAm | 15 | HEAm | 5 |
| 61 | F75-K25 | HEAm | 15 | NiPAm | 5 |
| 62 | H50-K50 | ALMP | 10 | NiPAm | 10 |
| 63 | H75-F25 | NiPAm | 15 | HEAm | 5 |
| 64 | I75-E25 | APTAC | 15 | HMAm | 5 |
| 65 | H75-E25 | ALMP | 15 | HMAm | 5 |
| 66 | A25-B75 | Am | 5 | DMAm | 15 |
| 67 | K50-D50 | NiPAm | 10 | MPAm | 10 |
| 68 | I100 | APTAC | 20 | - | 0 |
| 69 | E50-G50 | HMAm | 10 | tHMAm | 10 |
| 70 | E75-I25 | HMAm | 15 | APTAC | 5 |
| 71 | G75-F25 | tHMAm | 15 | HMAm | 5 |
| 72 | H25-K75 | ALMP | 5 | NiPAm | 15 |
| 73 | C75-D25 | DEAm | 15 | MPAm | 5 |
| 74 | A50-K50 | Am | 10 | NiPAm | 10 |

| | | | | | |
|---|---|---|---|---|---|
| 75 | H75-K25 | ALMP | 15 | NiPAm | 5 |
| 76 | C75-J25 | DEAm | 15 | AMPSAm | 5 |
| 77 | F100 | HEAm | 20 | - | 0 |
| 78 | A50-B50 | Am | 10 | DMAm | 10 |
| 79 | A50-D50 | Am | 10 | MPAm | 10 |
| 80 | D75-E25 | MPAm | 15 | HMAm | 5 |
| 81 | C75-A25 | DEAm | 15 | Am | 5 |
| 82 | I75-B25 | APTAC | 15 | DMAm | 5 |
| 83 | K75-F25 | NiPAm | 15 | HEAm | 5 |
| 84 | K50-F50 | NiPAm | 10 | HEAm | 10 |
| 85 | I75-C25 | APTAC | 15 | DEAm | 5 |
| 86 | D75-G25 | MPAm | 15 | tHMAm | 5 |
| 87 | H75-J25 | ALMP | 15 | AMPSAm | 5 |
| 88 | G75-E25 | tHMAm | 15 | HMAm | 5 |
| 89 | B50-F50 | DMAm | 10 | HEAm | 10 |
| 90 | J50-I50 | AMPSAm | 10 | APTAC | 10 |
| 91 | E75-F25 | HMAm | 15 | HEAm | 5 |
| 92 | F50-G50 | HEAm | 10 | tHMAm | 10 |
| 93 | K50-E50 | NiPAm | 10 | HMAm | 10 |
| 94 | E75-B25 | HMAm | 15 | DMAm | 5 |
| 95 | I25-J75 | APTAC | 5 | AMPSAm | 15 |
| 96 | C75-B25 | DEAm | 15 | DMAm | 5 |
| 97 | B75-F25 | DMAm | 15 | HEAm | 5 |
| 98 | J75-K25 | AMPSAm | 15 | NiPAm | 5 |
| 99 | B100 | DMAm | 20 | - | 0 |
| 100 | K50-G50 | NiPAm | 10 | tHMAm | 10 |
| 101 | C50-E50 | DEAm | 10 | HMAm | 10 |
| 102 | I75-F25 | APTAC | 15 | HEAm | 5 |
| 103 | A75-K25 | Am | 15 | NiPAm | 5 |
| 104 | A75-I25 | Am | 15 | APTAC | 5 |
| 105 | D75-C25 | MPAm | 15 | DEAm | 5 |
| 106 | K75-J25 | NiPAm | 15 | AMPSAm | 5 |
| 107 | I75-J25 | APTAC | 15 | AMPSAm | 5 |
| 108 | G50-C50 | tHMAm | 10 | DEAm | 10 |
| 109 | G75-C25 | tHMAm | 15 | DEAm | 5 |
| 110 | C75-E25 | DEAm | 15 | HMAm | 5 |
| 111 | K75-D25 | NiPAm | 15 | MPAm | 5 |
| 112 | A75-J25 | Am | 15 | AMPSAm | 5 |
| 113 | K50-I50 | NiPAm | 10 | APTAC | 10 |
| 114 | C100 | DEAm | 20 | - | 0 |

| | | | | | |
|---|---|---|---|---|---|
| 115 | G25-K75 | tHMAm | 5 | NiPAm | 15 |
| 116 | E75-G25 | HMAm | 15 | tHMAm | 5 |
| 117 | J75-E25 | AMPSAm | 15 | HMAm | 5 |
| 118 | I75-G25 | APTAC | 15 | tHMAm | 5 |
| 119 | G25-J75 | tHMAm | 5 | AMPSAm | 15 |
| 120 | B50-I50 | DMAm | 10 | APTAC | 10 |
| 121 | D50-I50 | MPAm | 10 | APTAC | 10 |
| 122 | F50-I50 | HEAm | 10 | APTAC | 10 |
| 123 | D50-G50 | MPAm | 10 | tHMAm | 10 |
| 124 | F50-E50 | HEAm | 10 | HMAm | 10 |
| 125 | B50-G50 | DMAm | 10 | tHMAm | 10 |
| 126 | J100 | AMPSAm | 20 | - | 0 |
| 127 | J50-C50 | AMPSAm | 10 | DEAm | 10 |
| 128 | J75-I25 | AMPSAm | 15 | APTAC | 5 |
| 129 | E100 | HMAm | 20 | - | 0 |
| 130 | A50-G50 | Am | 10 | tHMAm | 10 |
| 131 | H25-E75 | ALMP | 5 | HMAm | 15 |
| 132 | F75-E25 | HEAm | 15 | HMAm | 5 |
| 133 | H75-C25 | ALMP | 15 | DEAm | 5 |
| 134 | A50-F50 | Am | 10 | HEAm | 10 |
| 135 | D50-C50 | MPAm | 10 | DEAm | 10 |
| 136 | A50-J50 | Am | 10 | AMPSAm | 10 |
| 137 | B75-K25 | DMAm | 15 | NiPAm | 5 |
| 138 | H50-G50 | ALMP | 10 | tHMAm | 10 |
| 139 | C50-I50 | DEAm | 10 | APTAC | 10 |
| 140 | H25-C75 | ALMP | 5 | DEAm | 15 |
| 141 | B50-E50 | DMAm | 10 | HMAm | 10 |
| 142 | G50-I50 | tHMAm | 10 | APTAC | 10 |
| 143 | B75-E25 | DMAm | 15 | HMAm | 5 |
| 144 | H25-G75 | ALMP | 5 | tHMAm | 15 |
| 145 | F75-G25 | HEAm | 15 | tHMAm | 5 |
| 146 | K50-J50 | NiPAm | 10 | AMPSAm | 10 |
| 147 | K75-C25 | NiPAm | 15 | DEAm | 5 |
| 148 | F75-B25 | HEAm | 15 | DMAm | 5 |
| 149 | B75-I25 | DMAm | 15 | APTAC | 5 |
| 150 | H50-E50 | ALMP | 10 | HMAm | 10 |
| 151 | K100 | NiPAm | 20 | - | 0 |
| 152 | H50-C50 | ALMP | 10 | DEAm | 10 |
| 153 | J50-D50 | AMPSAm | 10 | MPAm | 10 |
| 154 | B75-D25 | DMAm | 15 | MPAm | 5 |

| | | | | | |
|---|---|---|---|---|---|
| 155 | B75-G25 | DMAm | 5 | tHMAm | 15 |
| 156 | H50-I50 | ALMP | 10 | APTAC | 10 |
| 157 | B75-J25 | DMAm | 15 | AMPSAM | 5 |
| 158 | E50-I50 | HMAm | 10 | APTAC | 10 |
| 159 | H75-G25 | ALMP | 15 | tHMAm | 5 |
| 160 | B50-D50 | DMAm | 10 | MPAm | 10 |
| 161 | B75-C25 | DMAm | 5 | DEAm | 15 |
| 162 | B50-C50 | DMAm | 10 | DEAm | 10 |
| 163 | H25-I75 | ALMP | 5 | APTAC | 15 |
| 164 | J75-C25 | AMPSAm | 15 | DEAm | 5 |
| 165 | J50-E50 | AMPSAm | 10 | HMAm | 10 |
| 166 | C75-I25 | DEAm | 15 | APTAC | 5 |
| 167 | F50-J50 | HEAm | 10 | AMPSAm | 10 |
| 168 | B50-J50 | DMAm | 10 | AMPSAm | 10 |
| 169 | F75-J25 | HEAm | 15 | AMPSAM | 5 |
| 170 | E25-K75 | HMAm | 5 | NiPAm | 15 |
| 171 | F75-I25 | HEAm | 15 | APTAC | 5 |
| 172 | D50-E50 | MPAm | 10 | HMAm | 10 |

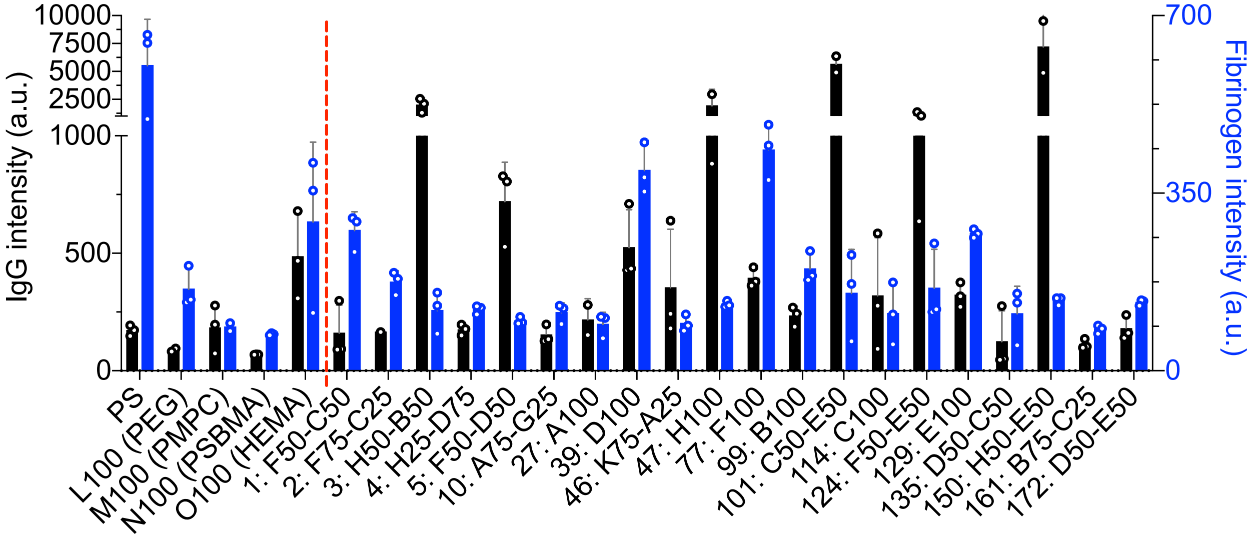

**Supplementary Fig. 2. IgG and fibrinogen adsorption assay.** IgG and fibrinogen adsorption on polystyrene (PS), gold standard and control hydrogels, and a selected subset of polyacrylamide hydrogels, detected with a fluorescence assay.

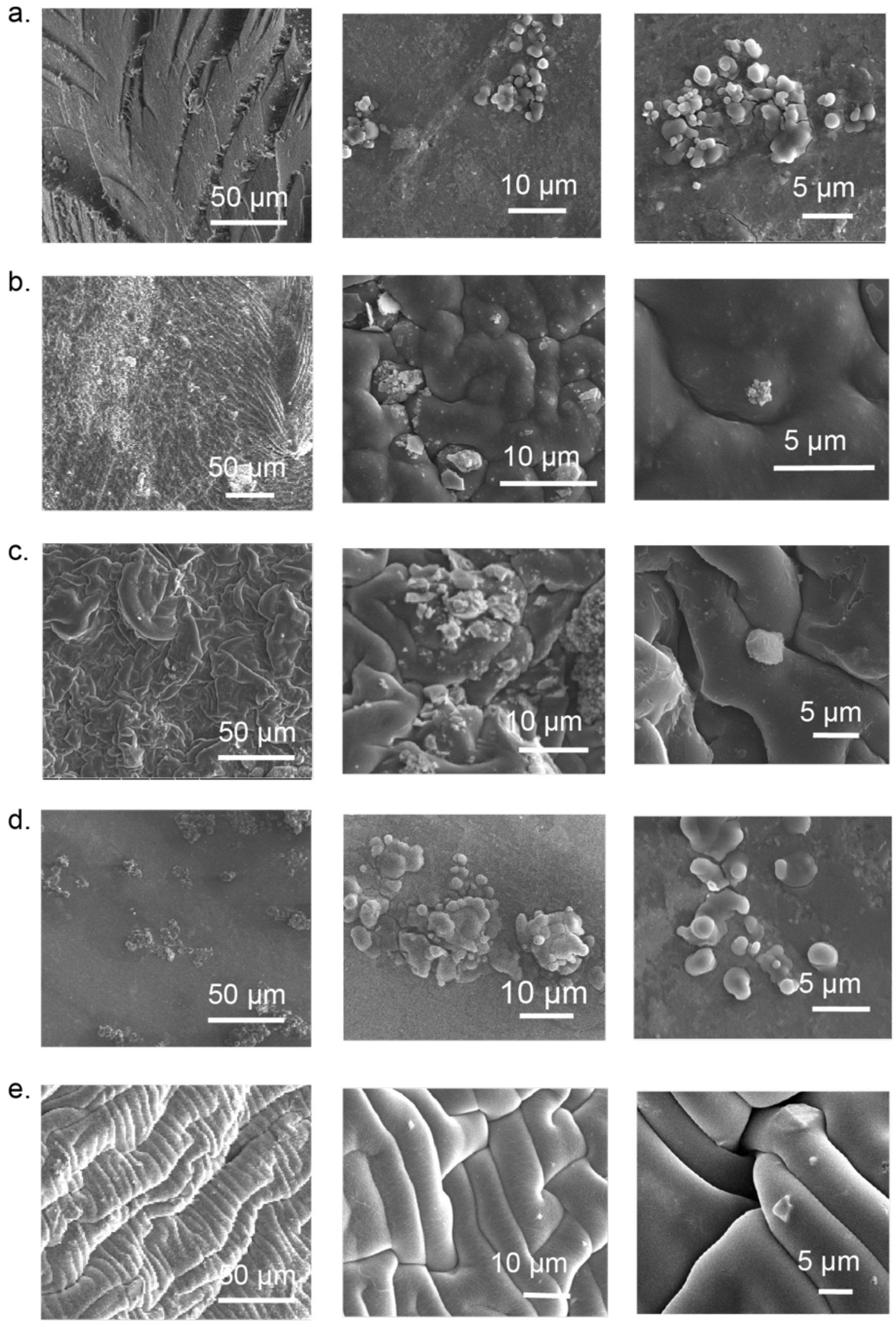

**Supplementary Fig. 3. SEM micrographs of adherent platelets on hydrogel samples.** SEM micrographs of hydrogel surfaces following 24 hr incubation with 50% serum and 1 hr incubation with platelet rich plasma. **a**. PEG. **b.** PMPC. **c.** PSBMA. **d.** PHEMA. **e.** F50-C50.

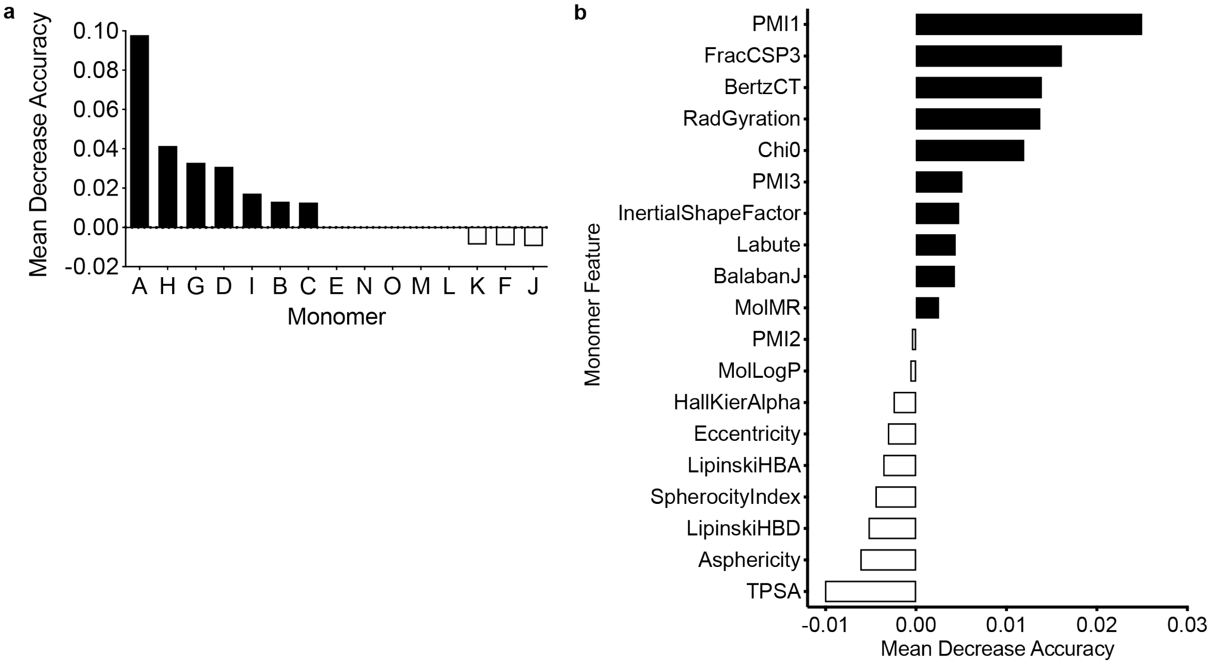

**Supplementary Fig. 4. Contribution of monomers and features in biofouling performance. a.** MDA feature relevance evaluated with feature set A that accounts for the monomer type and includes molar ratios as scaling factors. Monomers A, H, and G exhibited highest MDA, indicating high extent to which the feature is important for model development**. b.** Feature importance with Feature Set C. Positive MDA values indicate high relevance.

**Supplementary Table 3.** Performance of random forest classifier on the withheld test set with different features generated for the polymers.

*Feature Set A*

| | precision | recall | f1-score | support |
|---|---|---|---|---|
| 0 | 0.71 | 0.79 | 0.75 | 19 |
| 1 | 0.81 | 0.74 | 0.77 | 23 |
| micro avg | 0.76 | 0.76 | 0.76 | 42 |
| macro avg | 0.76 | 0.76 | 0.76 | 42 |
| weighted avg | 0.77 | 0.76 | 0.76 | 42 |

*Feature Set B*

| | precision | recall | f1-score | support |
|---|---|---|---|---|
| 0 | 0.71 | 0.79 | 0.75 | 19 |
| 1 | 0.81 | 0.74 | 0.77 | 23 |
| micro avg | 0.76 | 0.76 | 0.76 | 42 |
| macro avg | 0.76 | 0.76 | 0.76 | 42 |
| weighted avg | 0.77 | 0.76 | 0.76 | 42 |

*Feature Set C*

| | precision | recall | f1-score | support |
|---|---|---|---|---|
| 0 | 0.65 | 0.79 | 0.71 | 19 |
| 1 | 0.79 | 0.65 | 0.71 | 23 |
| micro avg | 0.71 | 0.71 | 0.71 | 42 |
| macro avg | 0.72 | 0.72 | 0.71 | 42 |
| weighted avg | 0.73 | 0.71 | 0.71 | 42 |

**Supplementary Table 4.** Definitions of molecular descriptors.

| Physical-Chemical Descriptors | | References |
|---|---|---|
| LabuteASA | *approximate surface area per Labute definition* | *J. Mol. Graph. Mod.* 18:464-77 (2000). |
| TPSA | *topological polar surface area* | *J. Med. Chem.* 43:3714-7, (2000) . |
| MolLogP | *partition coefficient, measure of lipophilicty* | Wildman and Crippen *JCICS* 39:868-73 (1999). |
| MolMR | *molar refractivity* | Wildman and Crippen *JCICS* 39:868-73 (1999). |
| Molecular Graph Descriptors | | |
| BalabanJ | *averaged distance sum connectivity, related to number of bonds between two vertices* | *Chem. Phys. Lett.* 89:399-404 (1982). |
| BertzCT | *index of molecular complexity due to distribution of the heteroatoms and complexity of the bonding* | *J. Am. Chem. Soc.* 103:3599-601 (1981). |
| Chi0 | *valence connectivity index* | *Rev. Comput. Chem.* 2:367-422 (1991). |
| HallKierAlpha | *describes number of atoms and number of connectivities* | *Rev. Comput. Chem.* 2:367-422 (1991). |
| H-bonding Descriptors | | |
| LipinskiHBA | *number of Ns and Os in the molecule* | - |
| LipinskiHBD | *number of N-H and O-H bonds in a molecule* | - |
| 3D Shape Descriptors | | |
| FracCSP3 | *number of tetrahedral carbon atoms* | - |
| InertialShapeFactor | *shape measured based on principle moments of inertia* | G. A. Arteca “Molecular Shape Descriptors” Reviews in Computational Chemistry vol 9. |
| PMI1 | *first (smallest) principle moment of inertia* | - |
| PMI2 | *second principle moment of inertia* | - |
| PMI3 | *third (largest) principle moment of inertia* | - |
| RadGyration | *radius of gyration* | G. A. Arteca “Molecular Shape Descriptors” Reviews in Computational Chemistry vol 9. |
| SpherocityIndex | *index of resemblance to that of a perfect sphere* | Todeschini and Consoni “Descriptors from Molecular Geometry” Handbook of Chemoinformatics. |
| Asphericity | *measurement of the shape of a refractive molecule and its effect on the bending of light* | A. Baumgaertner, “Shapes of flexible vesicles” J. Chem. Phys. 98:7496 (1993). |

| | | |
|---|---|---|
| Eccentricity | *measurement of the maximum distances from one atom vertex to any other atom vertex* | G. A. Arteca "Molecular Shape Descriptors" Reviews in Computational Chemistry vol 9. |

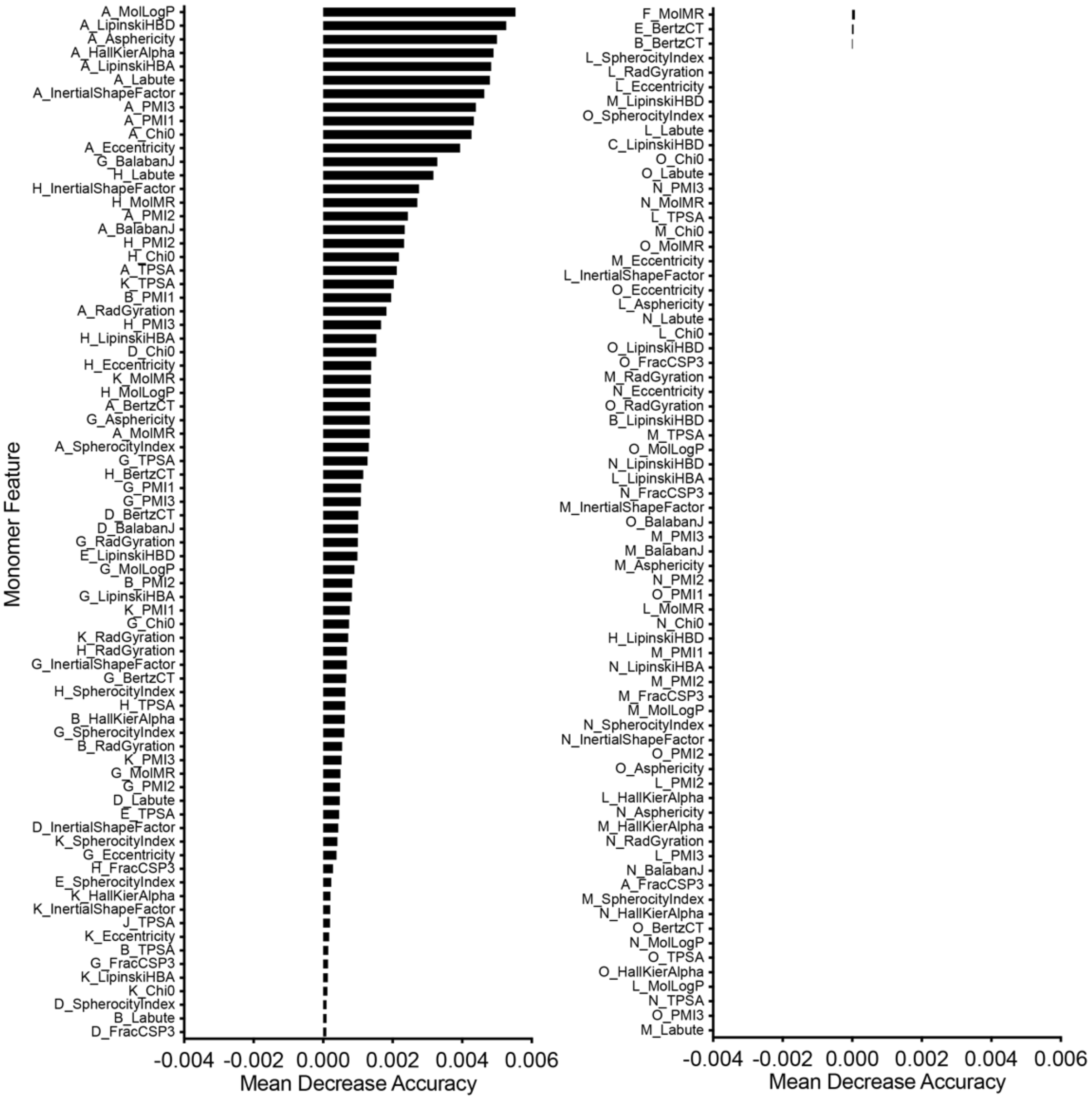

**Supplementary Fig. 5. Complete list of feature importance for each monomer from Feature Set B.** Positive MDA values indicate high impact of the feature (monomer-specific descriptor) to the model performance while negative MDA values indicate irrelevance of the feature.

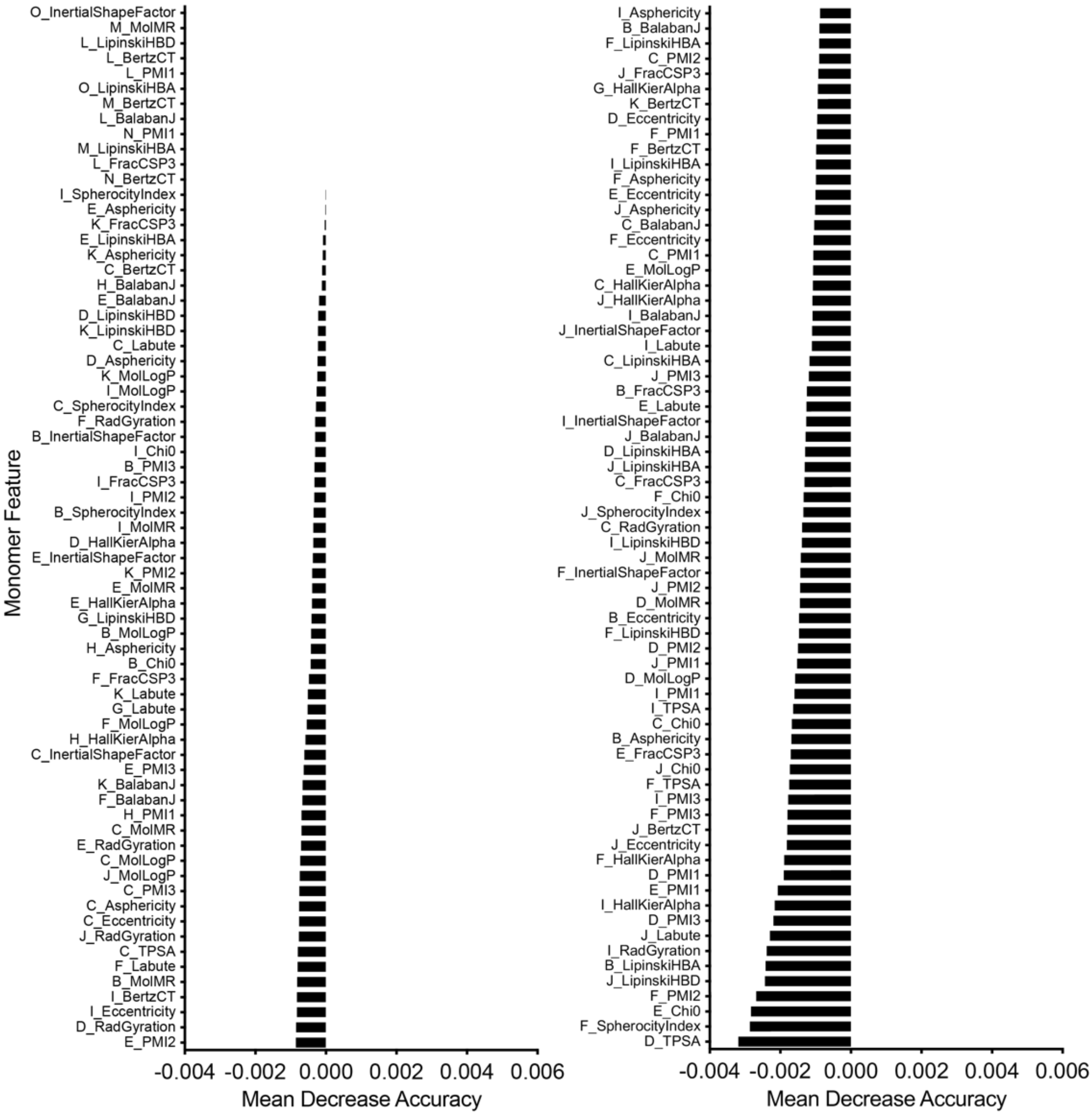

**Supplementary Fig. 5 (continued). Complete list of feature importance for each monomer from Feature Set B.** Positive MDA values indicate high impact of the feature (monomer-specific descriptor) to the model performance while negative MDA values indicate irrelevance of the feature.

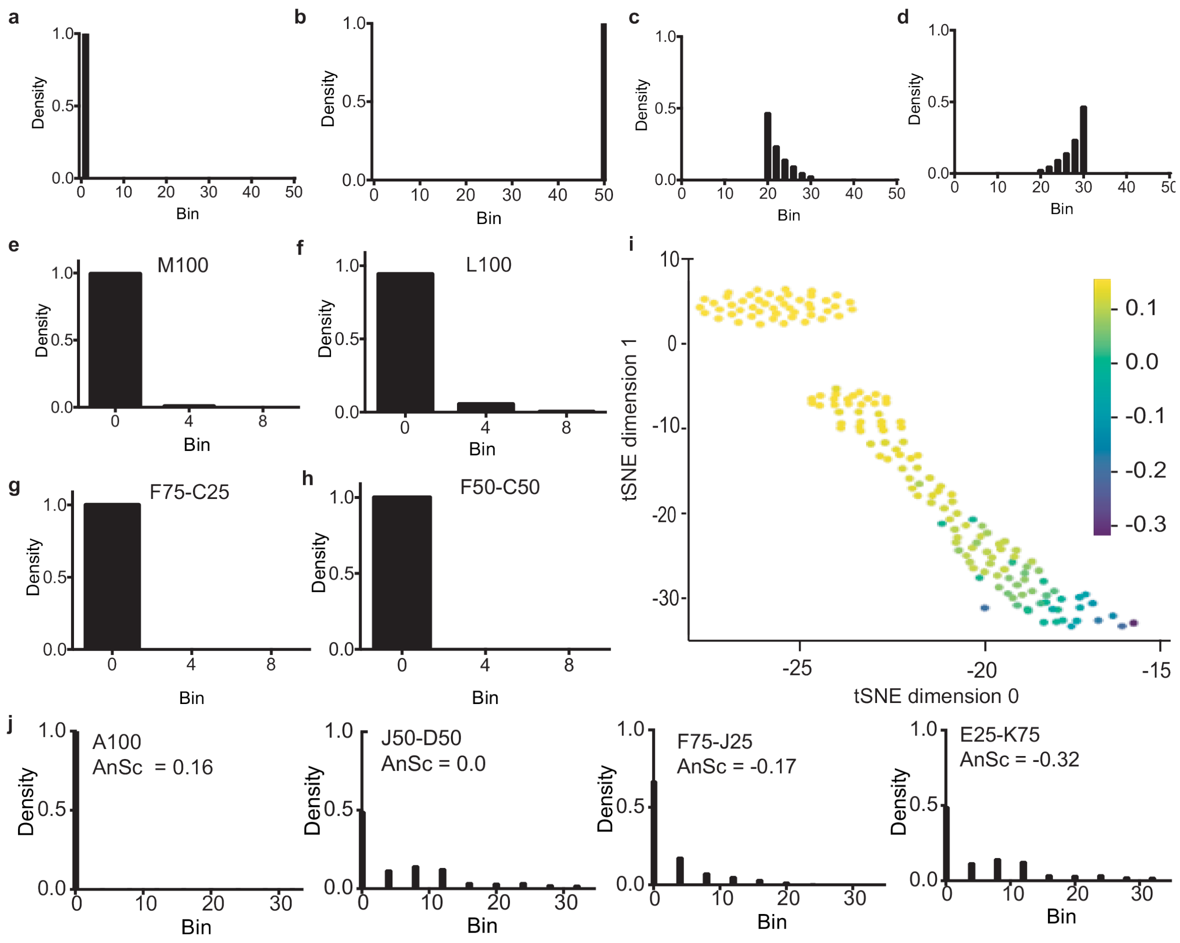

**Supplementary Fig. 6. Heterogeneity of platelet coverage suggests steric mode of fouling.** General types of population histograms that might be encountered: **a.** homogenous coverage, empty surface; **b.** homogenous coverage, uniformly covered surface; **c.** heterogenous coverage, covered surface with dense coverage domains. **d.** heterogenous coverage, covered surface with sparse domains. **e.** Population histogram of M100**. f.** Population histogram of L100. **g.** Population histogram of F75-C25**. h.** Population histogram of F50-C50. **i.** Results of the outlier detection in the manifold embedding of the population histograms via t-SNE algorithm. Color encodes the anomaly score of the sample obtained via Isolation Forest algorithm. Negative values of the anomaly score indicate outliers; high positive values indicate normal samples**. j.** Examples of population histograms, from the highest value (the group of the most normal samples, outlined by the box), to the lowest anomaly score (the most abnormal sample).

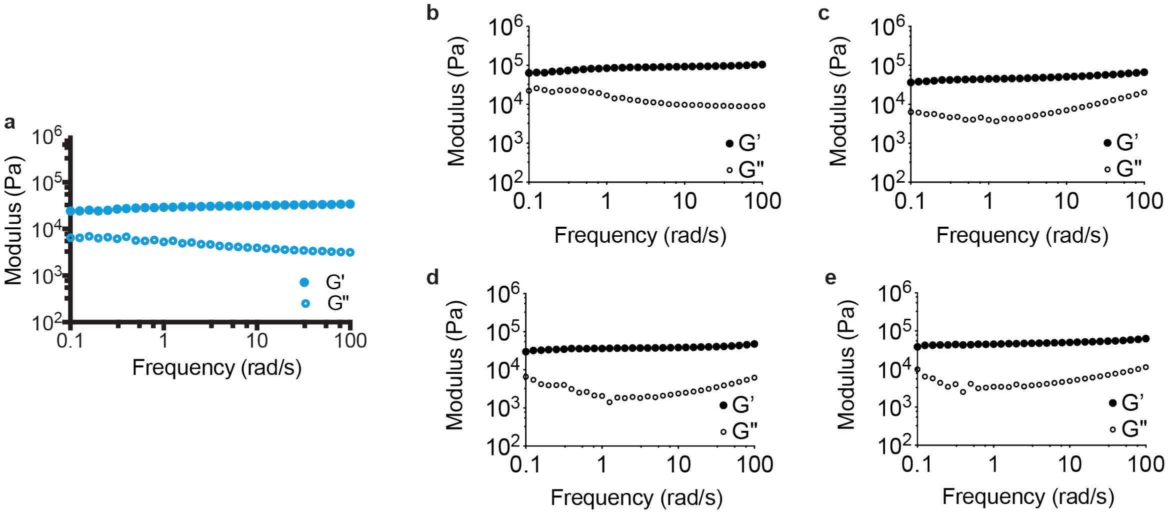

f

| | G’ (kPa) | G" (kPa) | tan(δ) | Mesh Size (nm) |
|---|---|---|---|---|
| F50-C50 | 27.0 ± 4.9 | 3.3 ± 2.5 | 0.12 ± 0.08 | 2.30 ± 0.10 |
| PEG | 66.7 ± 28.5 | 7.3 ± 4.6 | 0.13 ± 0.06 | 2.09 ± 0.03 |
| PMPC | 32.2 ± 9.4 | 4.9 ± 3.8 | 0.14 ± 0.09 | 2.85 ± 0.35 |
| PCBMA | 39.5 ± 12.8 | 8.4 ± 5.6 | 0.18 ± 0.08 | 2.93 ± 0.34 |
| HEMA | 41.8 ± 9.2 | 4.9 ± 0.3 | 0.12 ± 0.04 | 0.58 ± 0.25 |

**Supplementary Fig. 7. Rheological properties and mesh size of hydrogels.** Frequency-dependent rheological properties (1% strain) of hydrogels with low and high molecular weight monomers demonstrates that the resulting hydrogels exhibit only minor variations in observed *G’* and *G”* values. Oscillatory shear rheology was performed at 1% strain for **a.** F50-C50, **b.** PEG, **c.** PSBMA, **d.** PMPC, and **e.** PHEMA hydrogels. **f.** Summary of rheological properties and mesh size as determined by FRAP experiments.

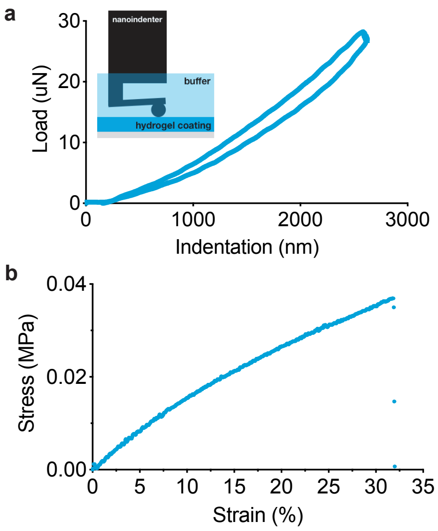

c

| Physical/Mechanical Property | Technique | Value (Mean ± S.D.) |
|---|---|---|
| G’ | Rheology | 27.0 ± 4.9 kPa |
| E | Nanoindentation | 1.0 ± 0.1 MPa |
| E | Tensile Test | 1.9 ± 0.2 kPa |
| Toughness | Tensile Test | 0.92 ± 0.42 J/m$^3$ |
| Strain at Break | Tensile Test | 35.1 ± 6.0 % |

**Supplementary Fig. 8. Mechanical and physical properties of F50-C50 hydrogel. a.** Representative curve of nanoindentation loading and unloading. **b.** Representative tensile test of F50-C50 hydrogel. **c.** Summary of properties determined from rheology, nanoindentation, and tensile testing.

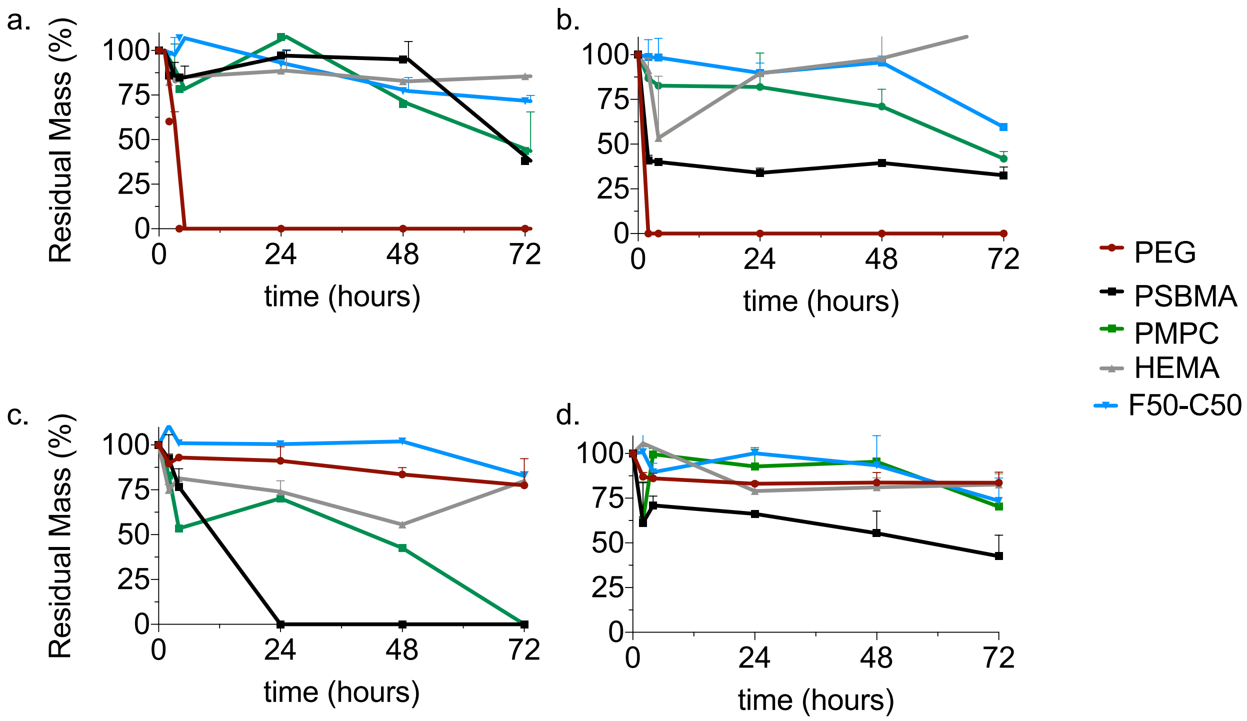

**Supplementary Fig. 9. Expedited degradation of hydrogels.** Expedited degradation of hydrogels was determined at 50 °C for gold standard materials, control hydrogels, and the top hydrogel formulation F50-C50. **a.** 12 M HCl. **b.** 12 M NaOH. **c.** 30% H2O2. **d.** 1 mg/mL lipase.

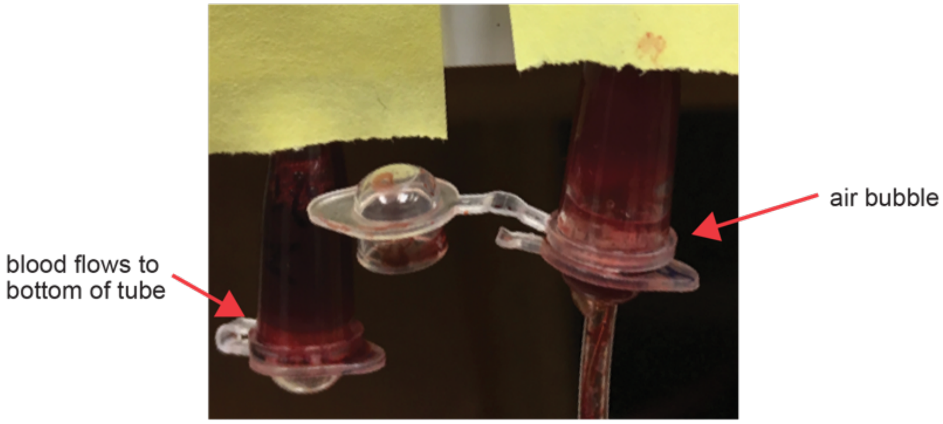

**Supplementary Fig. 10. Addition of calcium chloride expedites blood clotting in device testing.** Blood occlusion occurs completely after 6 hr in the presence of calcium chloride. With no $CaCl_2$ addition (left), blood flows freely, whereas with CaCl2 addition (right), blood samples clot and do not flow in the tube upon inversion.

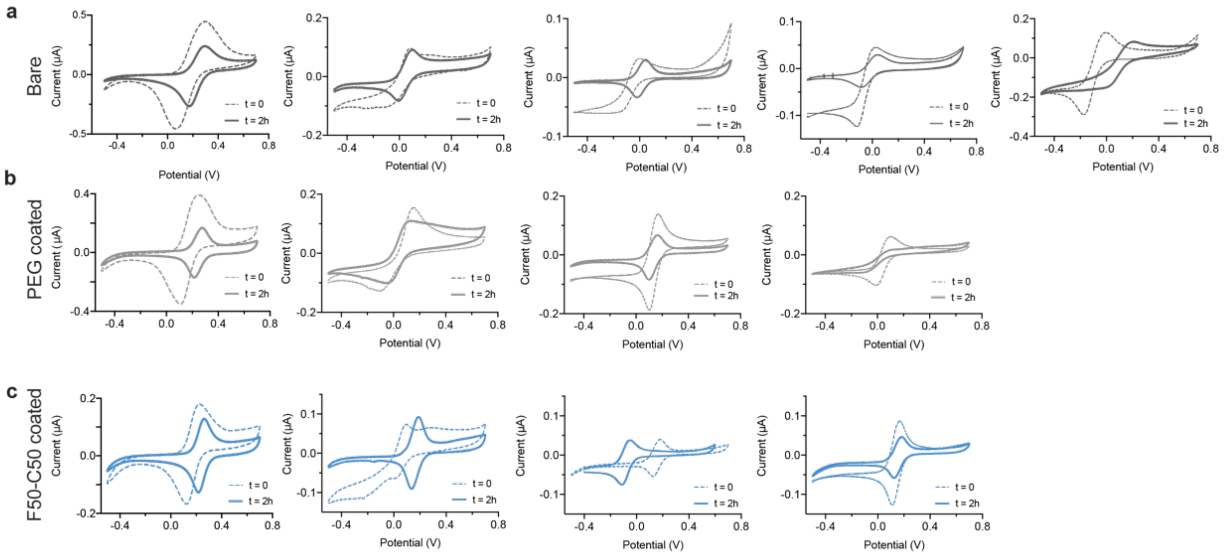

**Supplementary Fig. 11. Raw data for *in vitro* electrochemical device blood fouling assay**. Electrochemical device function in blood after 2 hr of incubation following CaCl2 addition. **a.** Bare device shows significant signal degradation. **b.** PEG-coated device. **c.** F50-C50-coated device.

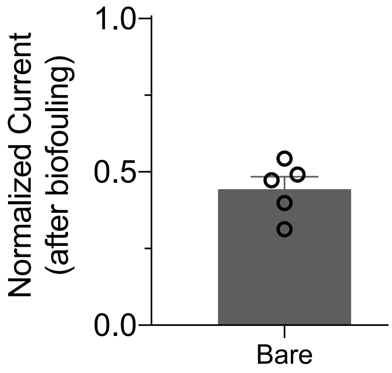

**Supplementary Fig. 12. Normalized signal intensity from blood fouling assay.** Normalized signal intensity, defined as the anodic peak current from CV, from bare devices after biofouling (after 2 hr with $CaCl_2$ addition). Bare probes (as described in the text) show severe signal degradation, with no statistical difference between PEG-coated and bare probes.

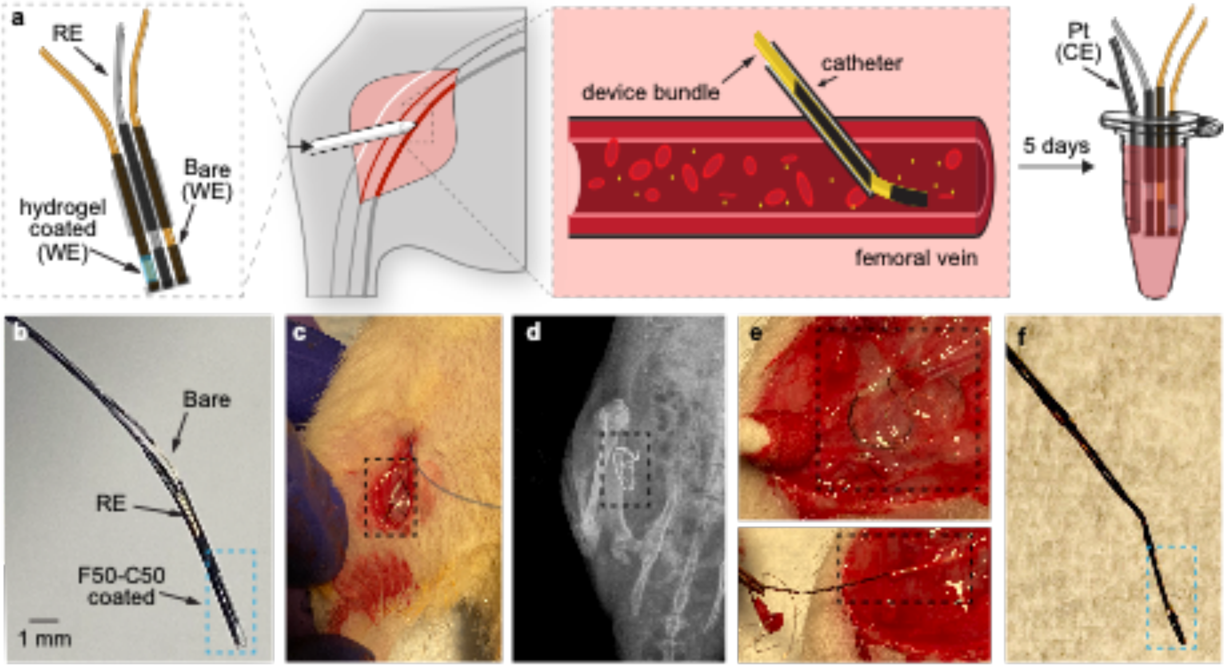

**Supplementary Fig. 13. Assessment of hydrogel-coated electrochemical sensors *in vivo*.** **a.** Bare and hydrogel-coated electrochemical probes were incubated in the femoral vein of a rat for 5 days prior to explantation and evaluation of device performance by CV. **b.** Image of electrochemical devices whereby the WE was coated with F50-C50 hydrogels. **c.** Probe insertion into the femoral vein of a rat. **d.** X-ray imaging indicates that the probe remains inserted in the femoral vein after 5 days. **e.** Fibrous encapsulation and removal of electrochemical probes after 5 days *in vivo*. **f.** Image after probe removal indicating that hydrogel coating on the devices remains intact.

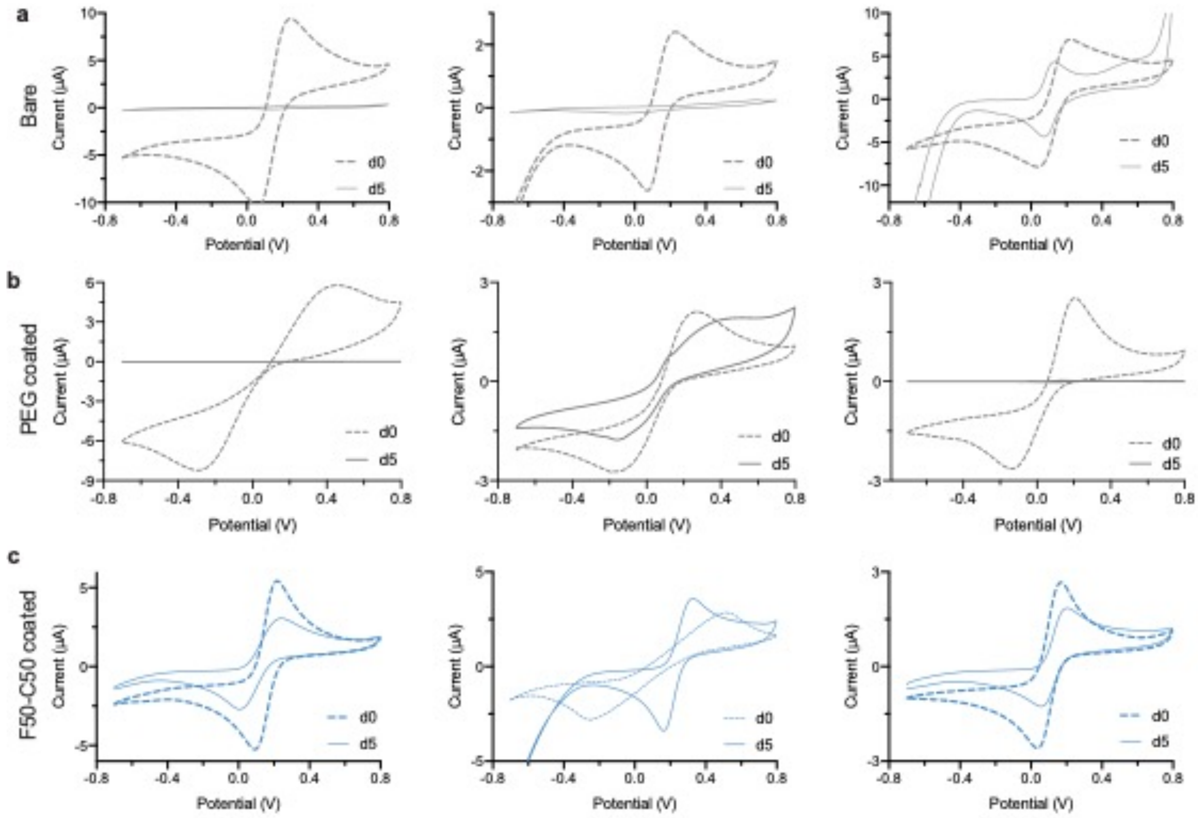

**Supplementary Fig. 14. Raw data for *in vivo* electrochemical device fouling assay.** Raw electrochemical sensor device data before and after 5 days of implantation in the femoral vein of a rat. **a.** Bare probe shows significant signal degradation. **b.** PEG-coated device. **c.** F50-C50-coated device.

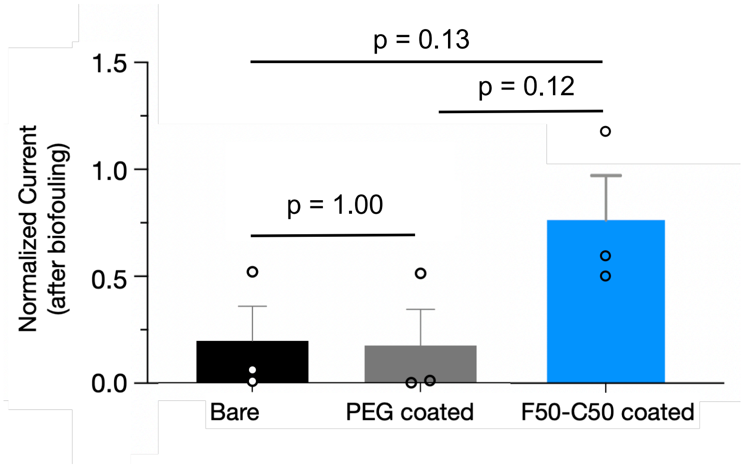

**Supplementary Fig. 15. Normalized signal intensity from *in vivo* fouling assay.** Normalized signal intensity, defined by anodic peak current from CV, of devices following 5 days of in vivo implantation in the femoral vein. This pilot study indicated that bare and PEG coated probes show dramatic signal degradation in comparison to F50-C50-coated probes. Only one probe from each of the three bare and PEG-coated probes tested showed any response, while all three F50-C50-coated probes showed robust signal. Data depicted as mean ± standard error with n = 3, and statistical analysis was conducted with a one-way ANOVA test with Tukey's multiple comparison test.

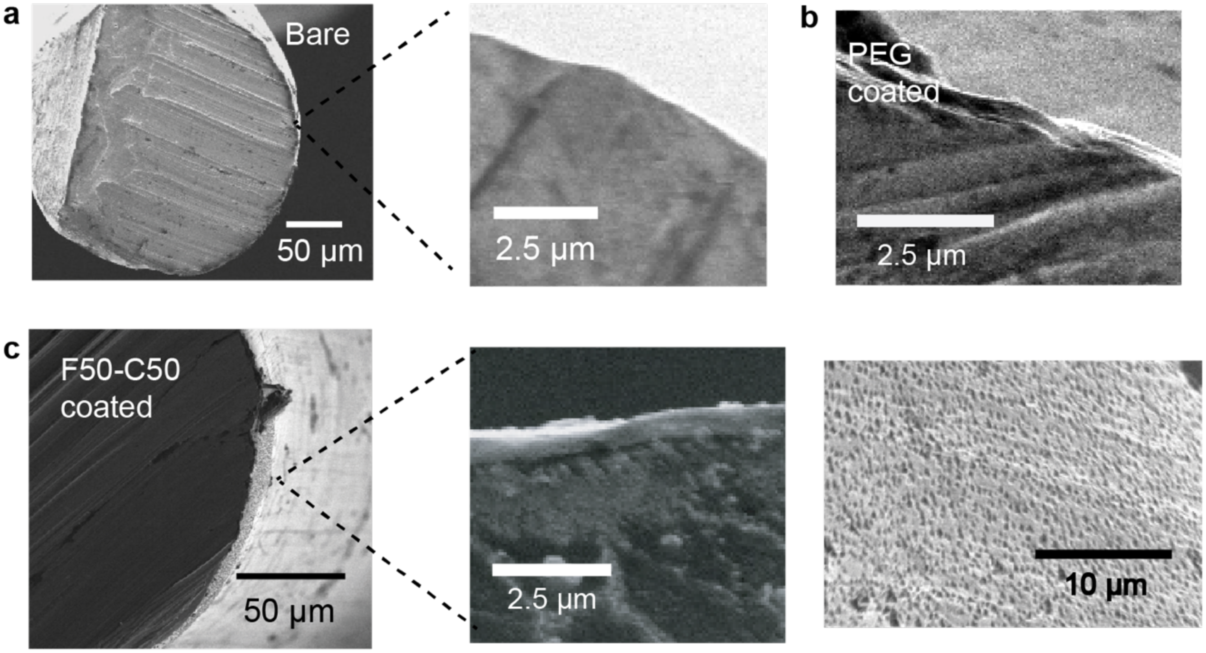

**Supplementary Fig. 16. SEM micrographs of aptamer probes. a.** Bare aptamer sensor**. b.** PEG-coated aptamer sensor. **c.** F50-C50 coated aptamer sensor.

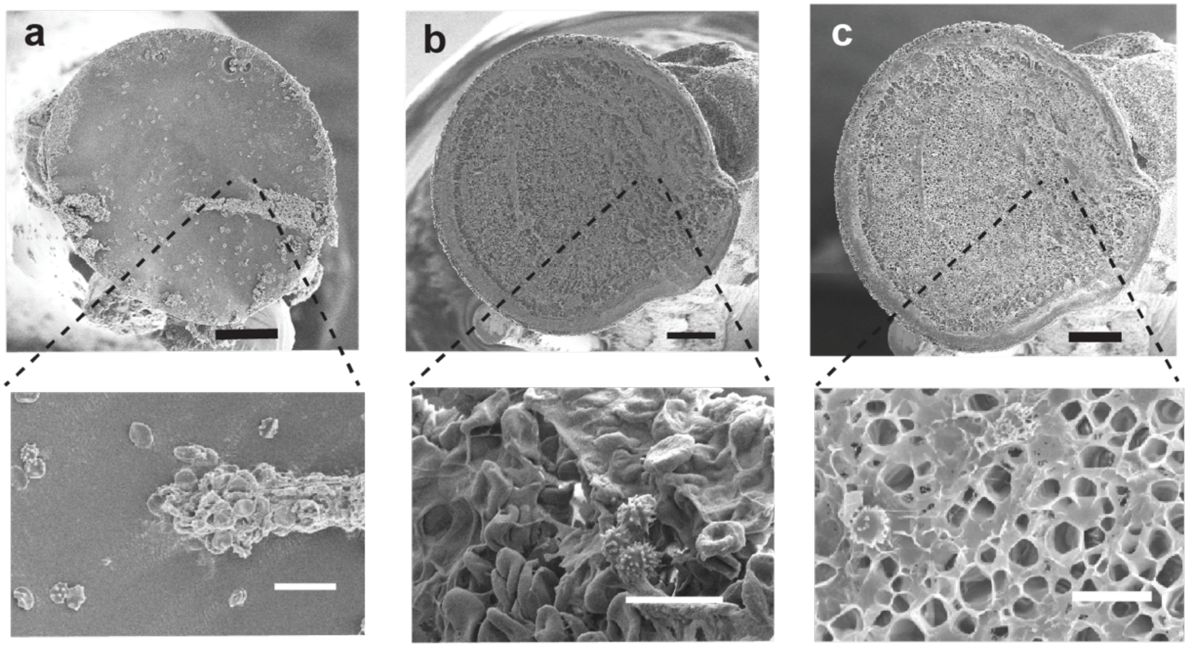

**Supplementary Fig. 17. SEM micrographs of DNA aptamer probes following blood fouling.** DNA aptamer probes after incubation in whole blood assay after 3 days. **a.** Platelet adhesion and activation on the tip of the bare probe. **b.** PEG-coated hydrogel shows coverage from platelet agglomeration. **c.** Homogenous coating of F50-C50 gel remains visible with less adhesion of platelets. Black bar = 100 µm. White bar = 10 µm.

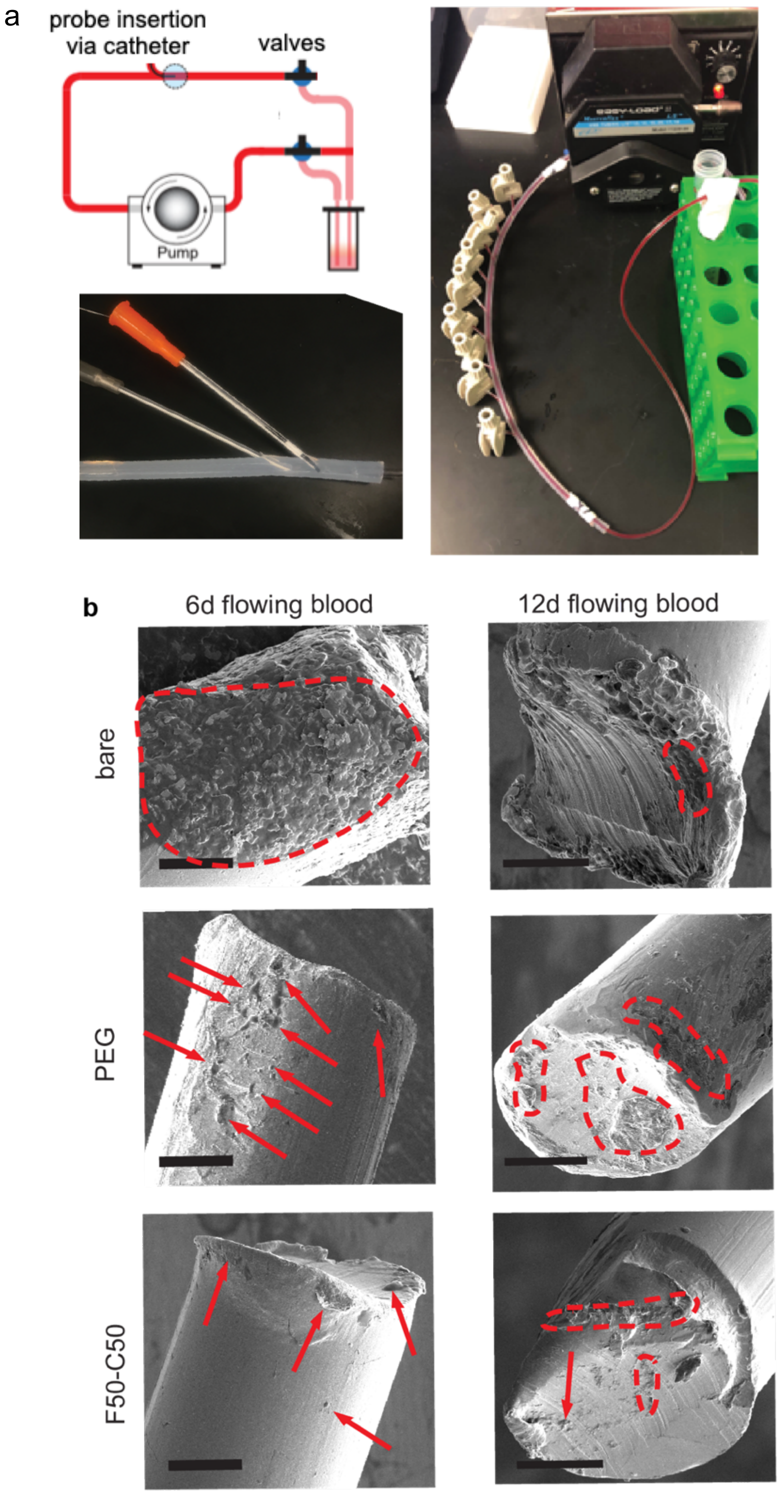

**Supplementary Fig. 18. DNA aptamer probes after flowing whole blood assay. a.** *in vitro* whole blood flowing assay. Probes are inserted through a catheter into tubing connected to pump head analogous to a closed loop blood pumping system**. b.** SEM Micrographs of the hydrogel coating on DNA aptamer probes, and the probe tips after 6 days and 12 days in flowing blood. Scale bar = 30 µm.

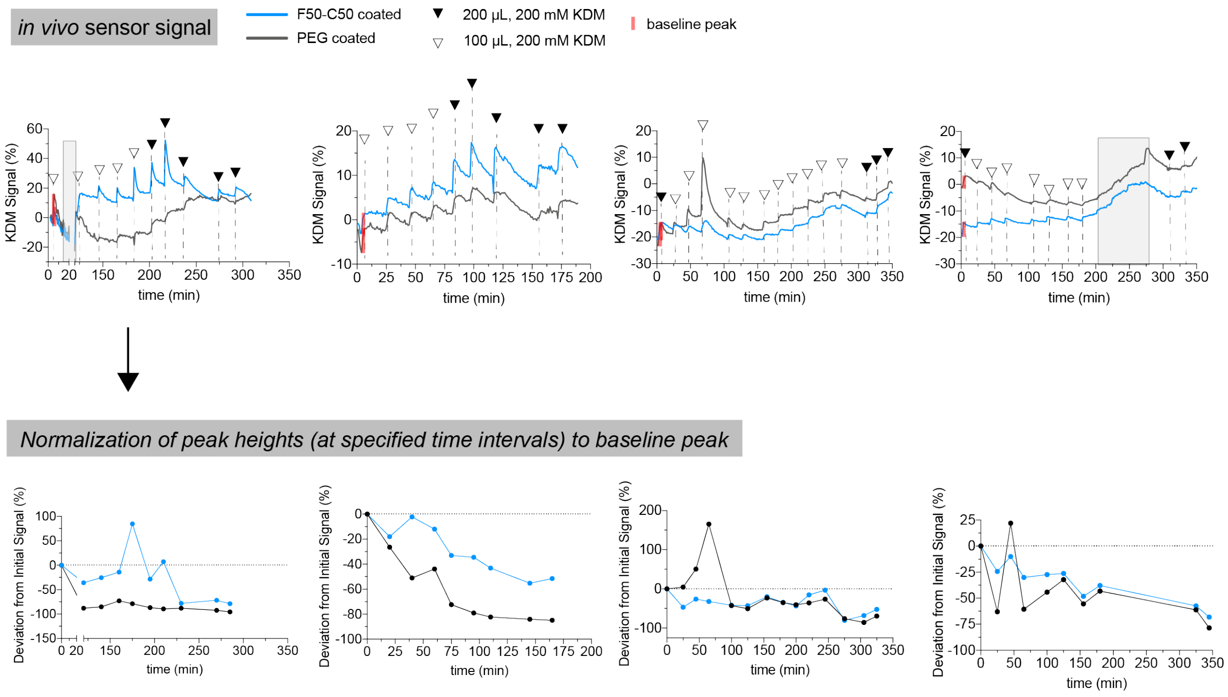

**Supplementary Fig. 19. Function of hydrogel-coated aptamer-based electrochemical sensors *in vivo*.** Application of PEG and F50-C50 hydrogels to DNA aptamer electrochemical sensors in vivo. (top) Raw data for real-time in vivo sensing of kanamycin. Filled arrows represent a 200 µL injection of 200 mM kanamycin and empty black arrows represent a 100 µL injection of 200 mM kanamycin. Shaded red boxes signal intensity at baseline after equilibration period of 1-2 hr. Gray box represents noise in data or signal acquisition was temporarily lost. (bottom) Peak heights were extracted from raw data at indicated time points and normalized to signal peak and dosage amount given at baseline time point.

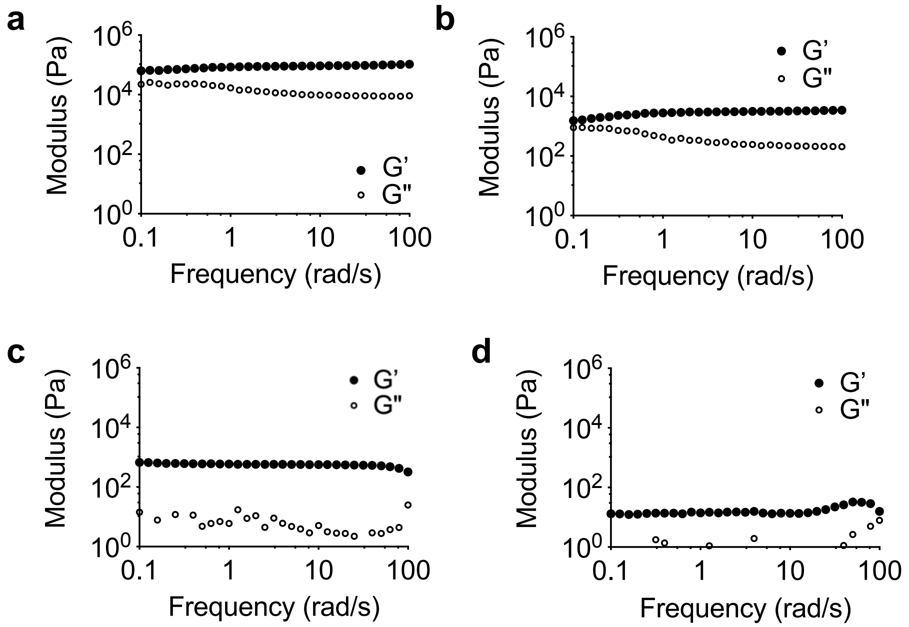

**Supplementary Fig. 20. Rheological characterization of alternative PEG hydrogels**. Several alternative PEG-based hydrogels were prepared comprising 20 wt% loading of PEG-based polymer. Frequency-dependent oscillatory shear rheology of each PEG hydrogel material performed at 1% strain: **a.** L100 (selected PEG control), **b.** PEGDMA740_PEGMA550, **c.** PEGDA10K, and **d.** PEGDMA10K.

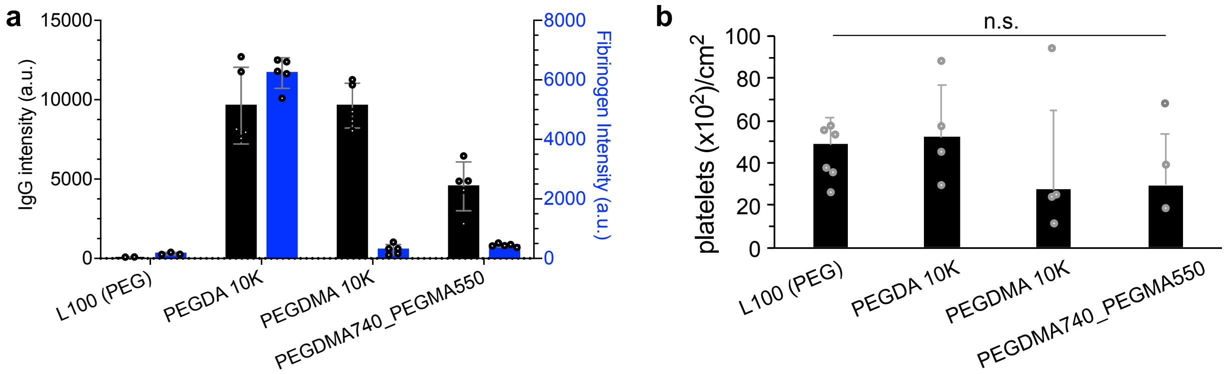

**Supplementary Fig. 21. Protein (IgG and fibrinogen) and platelet adhesion on alternative PEG hydrogels. a.** IgG and fibrinogen adsorption on PEG hydrogels. **b.** Platelet adhesion screening on PEG hydrogels. Median ± standard error platelet counts on hydrogel surfaces are shown with $n \geq 3$. Data was analyzed with a one-way ANOVA with Tukey's multiple comparison test where a threshold of significance was noted at $p = 0.05$.

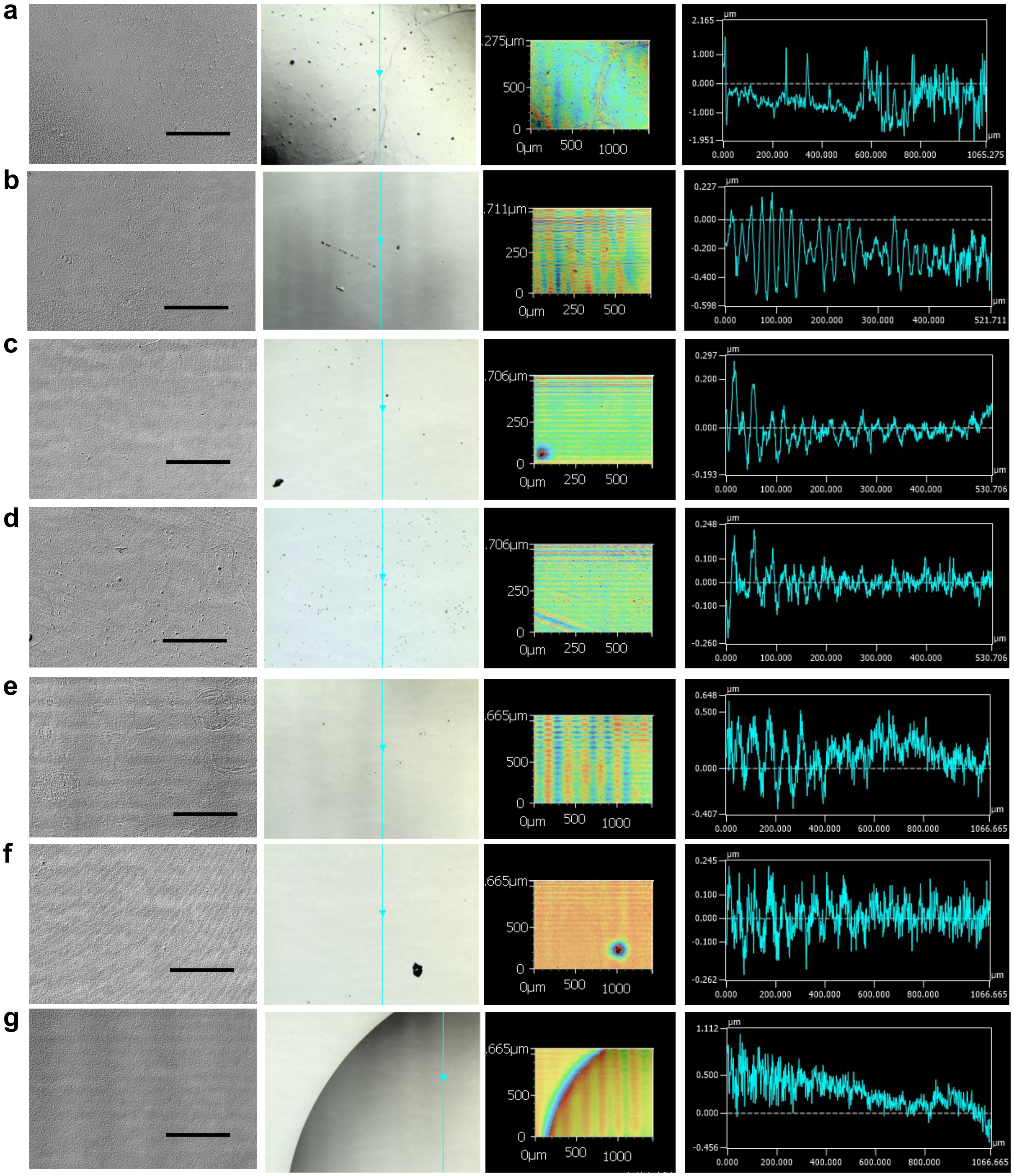

**Supplementary Fig. 22. Evaluation of surface roughness of hydrogels.** The arithmetical mean height of the surface features of PEG, the top performing PAAm hydrogel, homopolymer hydrogels comprising the top hydrogel, and a selected subset of PAAm gels are shown. Confocal images (10X; scale bar = 200 μm; column 1), overlaid laser and optical images (column 2), 3D profile (column 3), and profile graph (column 4) are shown for hydrogels: **a.** PEG, **b.** F50-C50, **c.** F100, **d.** C100, **e.** F75-A25, **f.** D75-C25, and **g.** D75-C25.

**Supplementary Table 5. Tabulated summary of arithmetical mean height of hydrogels**. Tabulated arithmetical mean height of the features of the hydrogel surfaces, defined as the absolute difference in height compared to the arithmetical mean of the surface, obtained from the experiments describe in Supplementary Fig. 21. Data represents mean ± s.d. with n = 3.

| Hydrogel formulation (platelet ranking) | Sa, arithmetical mean height (μm) |
|---|---|
| L100 (PEG) | 0.647 ± 0.55 |
| F50-C50 (1) | 0.966 ± 1.02 |
| F100 (77) | 0.100 ± 0.05 |
| C100 (114) | 0.310 ± 0.47 |
| F75-A25 (50) | 0.392 ± 0.30 |
| D75-C25 (105) | 0.283 ± 0.22 |
| B75-G25 (155) | 3.228 ± 2.61 |
